# An RNA to rule them all: Critical steps in Lassa virus ribonucleoparticle assembly and recruitment

**DOI:** 10.1101/2023.02.09.527830

**Authors:** Lennart Sänger, Harry M. Williams, Dingquan Yu, Dominik Vogel, Jan Kosinski, Maria Rosenthal, Charlotte Uetrecht

**Affiliations:** Bernhard Nocht Institute for Tropical Medicine, Bernhard-Nocht-Straße 74, 20359 Hamburg, Germany; CSSB Centre for Structural Systems Biology, Notkestraße 85, 22607 Hamburg, Germany; Leibniz Institute of Virology (LIV), Notkestraße 85, 22607 Hamburg, Germany; European Molecular Biology Laboratory Notkestraße 85, 22607, Hamburg, Germany; Structural and Computational Biology Unit, European Molecular Biology Laboratory, Meyerhofstraße 1, 69117 Heidelberg, Germany; Fraunhofer Institute for Translational Medicine and Pharmacology (ITMP), Discovery Research ScreeningPort, Schnackenburgallee 114, 22525, Hamburg, Germany; Faculty V: School of Life Sciences, University of Siegen, Am Eichenhang 50, 57076, Siegen, Germany; Deutsches Elektronen-Synchrotron (DESY), Notkestr. 85, 22607 Hamburg, Germany

## Abstract

Lassa virus is a negative-strand RNA virus with only four structural proteins that causes periodic outbreaks in West Africa. The nucleoprotein (NP) encapsidates the viral genome, forming the ribonucleoprotein complexes (RNPs) together with the viral RNA and the L protein. RNPs have to be continuously restructured during viral genome replication and transcription. The Z protein is important for membrane recruitment of RNPs, viral particle assembly and budding, and has also been shown to interact with the L protein. However, the interaction of NP, viral RNA and Z is poorly understood. Here, we characterize the interactions between Lassa virus NP, Z and RNA using structural mass spectrometry. We identify the presence of RNA as the driver for disassembly of ring-like NP trimers, a storage form, into monomers to subsequently form higher order RNA-bound NP assemblies. We locate the interaction site of Z and NP and demonstrate that while NP binds Z independently of the presence of RNA, this interaction is pH-dependent. These data improve our understanding of RNP assembly, recruitment and release in Lassa virus.

## Introduction

Lassa virus (LASV) is the causative agent of Lassa hemorrhagic fever and belongs to the *Arenaviridae* family within the *Bunyavirales* order. LASV is rodent-borne and endemic to West Africa infecting between 100,000-300,000 humans each year leading to approximately 5,000 deaths [1]. Recent predictions estimate up to 186 million people at risk of LASV infections in West Africa by 2030 [2]. Arenaviruses are bi-segmented negative-strand RNA viruses and each genome segment encodes for two structural proteins using an ambisense coding strategy. The large genome segment encodes for the ∼250 kDa large (L) protein, which contains the RNA-dependent RNA polymerase, and a small RING finger matrix protein Z, which has a molecular weight of ∼ 11 kDa. The small genome segment encodes for the viral glycoprotein precursor (GPC, ∼75 kDa) and the nucleoprotein (NP, ∼63 kDa) [3].

Despite the central role of the LASV L protein in viral genome replication and transcription, NP is also essential for these processes [4]. It encapsidates the viral RNA genome forming – together with L – the ribonucleoprotein complex (RNP). Within the RNP the viral RNA is protected from degradation and recognition by cellular pattern recognition receptors. The RNP is the structural and functional unit of viral genome replication and transcription [5]. NP is composed of an N-terminal and C-terminal domain. The N-terminal domain contains an RNA binding cavity and the C-terminal domain an exonuclease activity [6, 7]. When not bound to RNA, NP forms homotrimeric rings with a head to tail arrangement of the NP molecules. In this conformation, RNA binding is hindered as the respective pocket is blocked by two helices [8]. To allow for RNA binding an RNA gating mechanism was proposed where the C-terminal domain rotates slightly away allowing one helix to become partly unstructured (α5) whereas the other helix (α6) shifts away from the RNA binding groove [9]. However, the trigger and mechanism of this conformational change are unknown.

The matrix protein Z lines the inner side of the viral envelope and mediates virion assembly and budding by homo-oligomerization and interaction with host factors [10-12]. It is anchored into membranes by N-terminal myristoylation [13] and has been hypothesized to mediate recruitment of RNPs to the plasma membrane for virion assembly by direct or indirect interaction with NP [10, 14, 15]. Z has also been shown to interact with the L protein inhibiting its polymerase activity potentially by blocking the formation of an elongation conformation of L [16-21]. It remains unclear whether RNP recruitment is driven by the interaction between NP and Z, L and Z and/or via a host protein.

In this study we investigate the mechanism and dynamics of RNP formation and recruitment. In presence of RNA, even short ones, the ring-like NP trimer undergoes conformational changes leading to its disassembly into RNA bound NP monomers, which subsequently form higher order NP-RNA assemblies, reminiscent of RNPs. Additionally, we show that LASV NP can directly interact with the Z protein via the C-terminal domain. We demonstrate that this interaction is strongly reduced at low pH, which would be in line with the conditions during virus entry via the endosomal pathway to initiate the release of RNPs from the matrix.

## Results

The full-length LASV NP and Z proteins (AV strain) were expressed in *E. coli* and purified for subsequent interaction studies. As described previously, both NP and Z can form oligomers [22, 23]. While NP eluted from the size exclusion column (SEC) in one peak corresponding to a trimer, Z eluted in two peaks indicating a monomeric and oligomeric state. The oligomerization of the proteins was further analyzed by native mass spectrometry (nMS) [24]. We found that NP was mainly trimeric (189.56 +/-0.03 kDa), which will be referred to as NP_3_, but also traces of NP hexamers could be detected. The hexameric NP was already found by others with a retained exonuclease function [25]. However, it is unclear whether the hexamer is present during viral infection inside cells (or just an artifact from in vitro expression) nor what its biological role would be. Further characterizations of NP_3_ using nMS revealed extraordinarily high gas phase stability. No collision-induced dissociation (CID) was observed at up to 200 V acceleration into the collision cell (Fig. 1). Beyond 200 V, fragmentation into peptides took place but no monomer dissociation, pointing towards a high non-covalent complex stability in the gas phase in the absence of disulfide bonds. Although the hexameric NP is not the focus of this study, it is worth mentioning that it also displays a high gas phase stability. Even at 300 V acceleration voltage, the hexameric NP remained stable pointing to a specific and strong interaction. For the Z protein, one of the two fractions from SEC could be identified as monomeric Z (11.35 +/-0.04 kDa), whereas the second fraction contained heterogeneous oligomers of Z (Fig. S1) in the range of around 120 kDa. All following experiments were performed using the monomeric fraction of Z.

**Figure 1:**
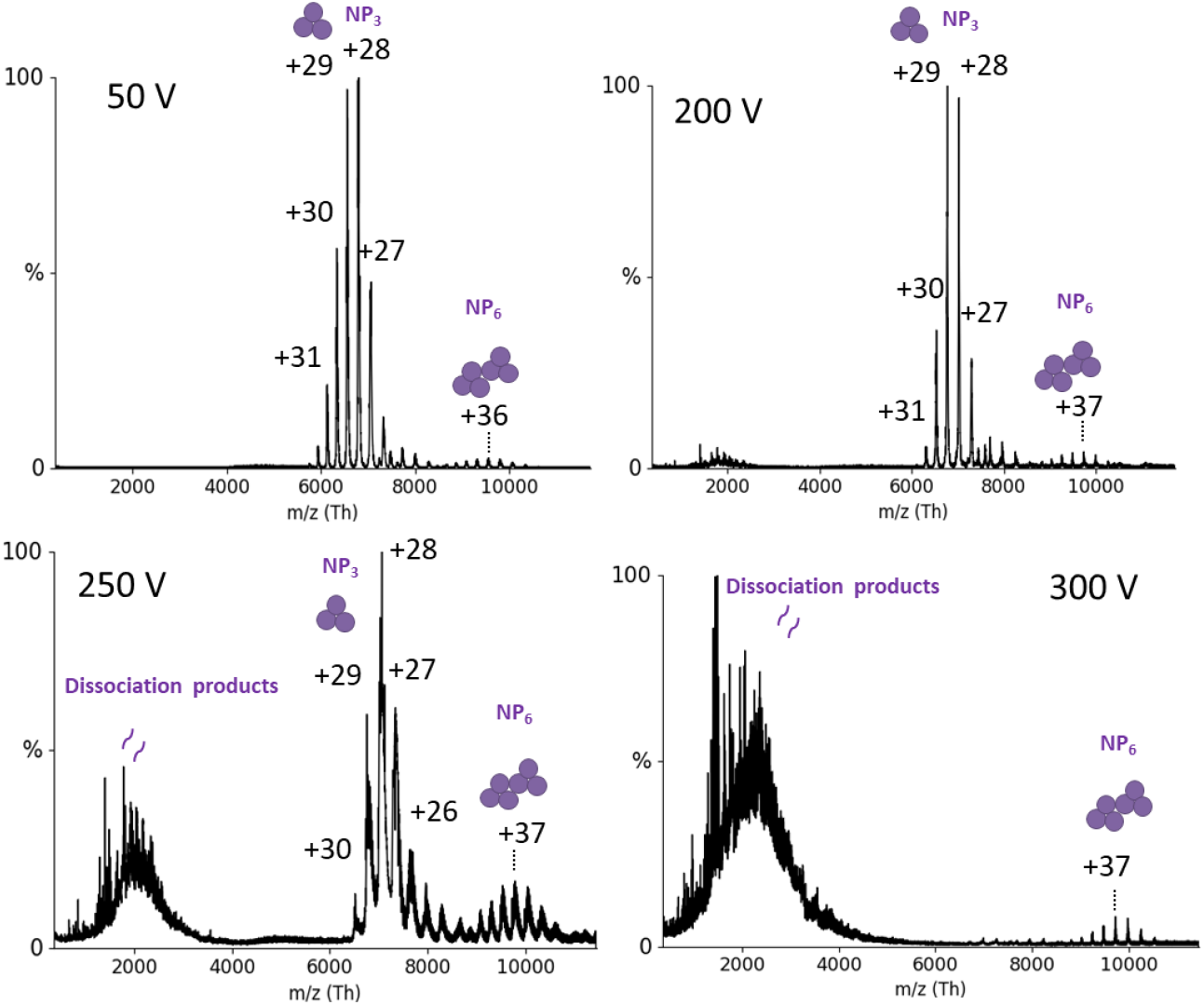
nMS of NP at different acceleration voltages (V). 6 μM of NP was measured in 150 mM ammonium acetate buffer surrogate at pH 7.5 with acceleration voltages in the collision cell between 50 V and 300 V (indicated in the graphs). The mass of the species between *m/z* 6,000-8,000 was determined as 189.56 +/-0.03 kDa which corresponds to an NP trimer. Charge states are indicated. Around *m/z* 10,000 an NP hexamer is detectable and above 200 V peptide fragments appear in the low mass range.

### NP directly interacts with monomeric Z protein

It was previously shown that LASV NP colocalizes with Z in cells [10]. We analyzed the ability of NP and Z to form complexes by direct interaction using purified proteins. nMS measurements of the trimeric NP with a 10-fold excess of Z showed new peaks appearing in the mass-to-charge (*m/z*) range between 5,000 and 7,000 (Fig. 2 A-C). To analyze the exact complex composition in this area, we used tandem MS (MS/MS) with CID. CID-MS/MS of peaks corresponding to 1, 2 or 3 Z proteins attached to an NP_3_ resulted in dissociation products of NP_3_ (189.56 +/-0.03 kDa), NP_3_ + Z (201.00 +/-0.09 kDa) and NP_3_ + 2xZ (212.37 +/-0.13 kDa), respectively (Fig. 2D).

**Figure 2:**
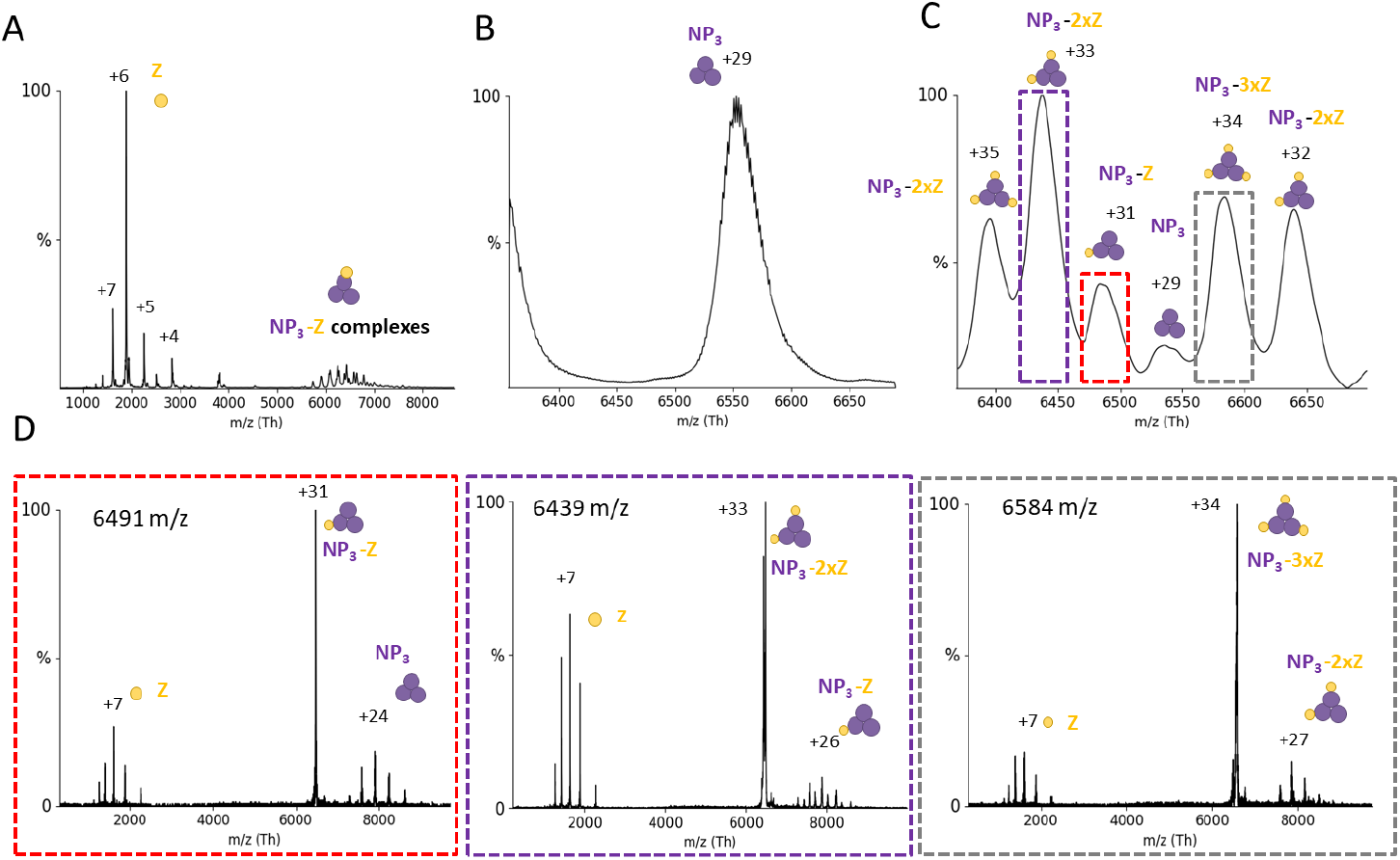
NP_3_ binds up to 3 Z monomers: NP and Z were incubated in a 1:10 molar ratio and subjected to nMS in a 150 mM ammonium acetate solution at pH 7.5. (A) Overview spectrum recorded with 50 V acceleration. Two peak series are observable between 1,000 and 4,000 *m/z* (corresponding to the Z protein) and between 5,500 and 7,000 *m/z* (NP_3_ complexes). Zoom to NP_3_ 29+ charge state without (B) and with Z protein (C). Several new species corresponding to NP-Z complexes are recorded. (D) CID-MS/MS at 100 V acceleration for NP_3_-Z (31+ charge state), NP3-2xZ (33+ charge state) and NP_3_-3xZ (34+ charge state). CID products correspond to masses for Z (11.35 +/-0.04 kDa), NP_3_ (189.56 +/-0.03 kDa), NP_3_-Z (201.00 +/-0.09 kDa) and NP3-2xZ (212.37 +/-0.13 kDa), respectively. The charge states of the most abundant peaks are indicated.

Overall, these experiments demonstrate a direct interaction between LASV NP_3_ and Z, in an RNA-free context with a stoichiometry of one binding interface per NP monomer. To determine the affinity of Z for NP_3_, we recorded nMS spectra in triplicates at an NP to Z molar ratio of 1:3 and used the deconvoluted spectra to determine the area under the curve (AUC). We determined an apparent K_D_ of 110 μM (± 10) for binding of a Z protein to an NP_3_ (Fig. S2).

Next, we tested whether NP-Z interaction depends on the oligomerization state of NP using an NP trimerization-deficient mutant (R52A). Elution profile of the SEC for NP-R52A showed two peaks corresponding to a trimeric and monomeric state. We analyzed both fractions by nMS and found additional traces of dimers in the monomeric fraction. (Fig. 3A, S3). The monomeric fraction was used for all following experiments. Interestingly, we observed heterogenous mass species with a mass difference of always around 300 Da (Fig. S3). This indicates different numbers of nucleotide monophosphates (NMPs) from the expression host being bound by the NP-R52A. These NMPs can be displaced when adding RNA to NP-R52A. At a 1:3 ratio between NP-R52A and Z, native mass spectra clearly showed the 75.5 +/-0.3 kDa monomeric NP-R52A-Z complex (Fig. 3B) with a K_D_ of 33 (± 2) μM. CID-MS/MS of the NP-R52A-Z complex further confirmed this interaction (Fig. S3). These data show that the monomeric NP-Z interaction is stronger compared to trimeric NP, either influenced by the monomeric conformation or by the arginine mutation.

**Figure 3:**
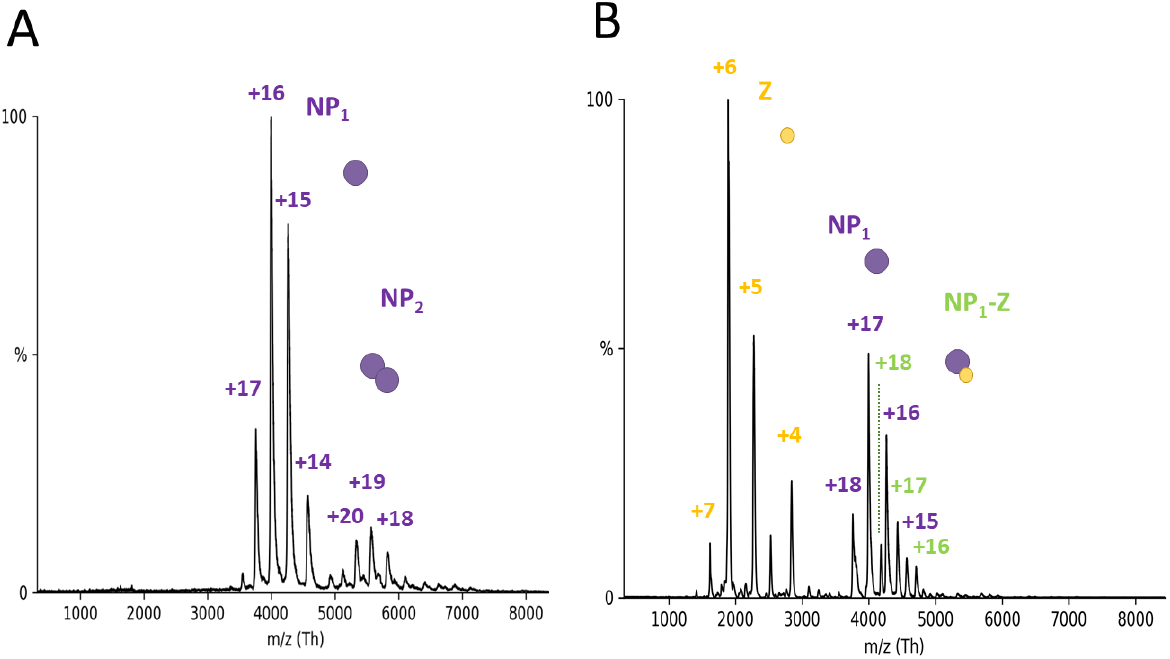
NP-Z interaction is independent of the oligomerization state of NP in nMS. (A) NP trimerization mutant R52A was measured with nMS in a 150 mM ammonium acetate solution at pH 7.5 and a concentration of 5.2 μM. Acceleration voltage in the collision cell was at 50 V for the depicted spectrum. R52A appears primarily in the monomeric state between 3,500 and 4,500 with charge states between +17 and +14 and a corresponding mass ofM 63.90 +/-0.08 kDa. A small fraction of dimeric NP is visible in the *m/z* range between 5,000 and 6,000 with the corresponding charge state of +20, +19 and +18. The most abundant species was normalized to 1. (B) R52A and Z were incubated in a 1:3 ratio and measured with the same instrument settings as described in A. Spectra show new peak series appearing for unbound Z protein in the *m/z* range between 1,500 and 3,000. New peaks also appear in the range between 4,000 and 5,000 *m/z* with +18, +17 and +16 charge states corresponding to a complexation of R52A and Z (75.5 +/-0.3 kDa).

We then used an integrative structural approach to map the interaction site between NP and Z. First, we predicted the structure of NP-Z complex by AlphaPulldown [26], a python package built upon AlphaFold [27] and AlphaFold Multimer [28] (Fig. 4A-B). The structural model obtained high local and global quality scores as returned by AlphaFold and AlphaPulldown (Fig. S4, Tab. S1) and predicted that Z interacts with the C-terminal domain of NP (Fig. 4B). To map changes in the protein dynamics caused by protein-protein interactions, we used hydrogen deuterium exchange MS (HDX-MS)[29]. This method is complementary to nMS and can be carried out in standard buffers. We used this method to experimentally validate the predicted interaction site. Labeling experiments were performed with NP both in the presence and absence of Z protein. Differences in deuterium uptake between both states were mapped to the NP sequence (Fig S5, S6). We found significant differences in deuterium uptake for the peptides 450-483 and 456-483, which align with the predicted NP-Z interface (Fig. 4C). The differences based on the deuterium uptake at the 6h timepoint were plotted to the predicted NP-Z complex (Fig. 4D, S6).

**Figure 4:**
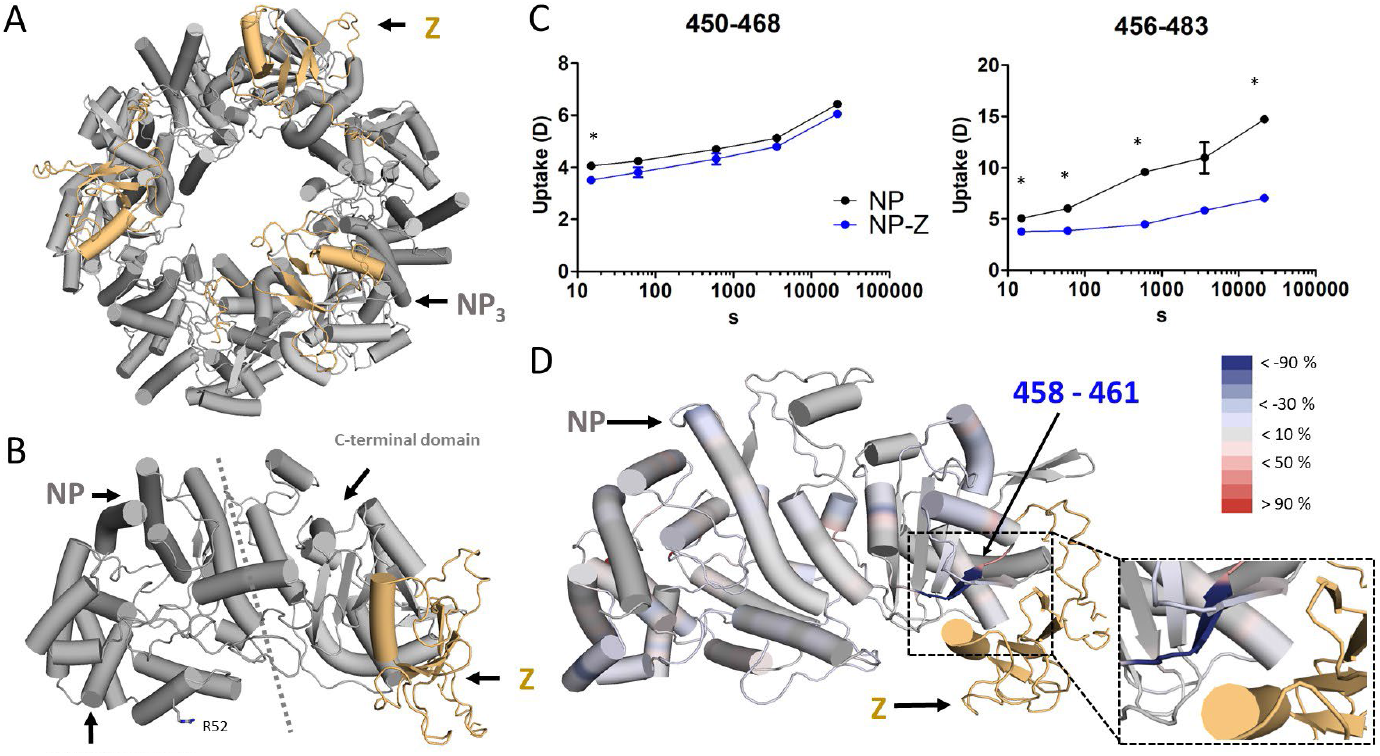
Identification of the NP-Z interface. The trimeric model of NP-Z (A) was generated by superposing the model of the NP-Z dimer predicted by AlphaFold (Version 2.1.0) (B) onto the crystal structure of the NP trimer (PDB ID 3R3L). NP si colored grey and the Z protein is shown in gold. (C) HDX differences in NP peptides in presence of Z. Uptake plots comparing the conditions of NP alone and NP-Z mixture with significant differences (*delta HDX > X D – 95 % CI, see methods, Table S2) around the predicted NP-Z interaction site (D). Differences in deuterium uptake between both states were mapped to the NP sequence and can be seen in Fig. S6. The differences are based on the atomic range which considers overlapping peptides with no differences. The differences at the 6h timepoint were plotted onto the predicted NP-Z complex. Regions that are protected and exposed in HDX in the presence of Z are highlighted in blue and red, respectively. Regions in grey show no change in deuterium uptake in the presence of Z. The Z protein is highlighted in gold.

Next, we introduced point mutations into the Z protein based on the NP-Z structural model to further validate the proposed interface. We exchanged polar residues to alanine or glycine at or in close proximity to the predicted interface. These were R4, R16, S59, S61, N62, R74, and T82. We measured all Z mutants together with NP in a 1:3 molar ratio by nMS and compared NP-Z complex formation by using the deconvoluted spectra of at least 3 independent measurements (Fig. 5A). Z mutants R16A and R74A showed strong reduction of NP-Z complex formation compared to the wild-type Z, whereas the other mutations had no effect on NP-Z interaction (Fig. 5B). This leads to the conclusion that NP-Z interaction is strongly dependent on arginine 16 and 74 of Z, which supports the predicted model (Fig. 5C). The other point mutations of polar residues seem to be less critical and were not able to influence NP-Z interaction.

**Figure 5:**
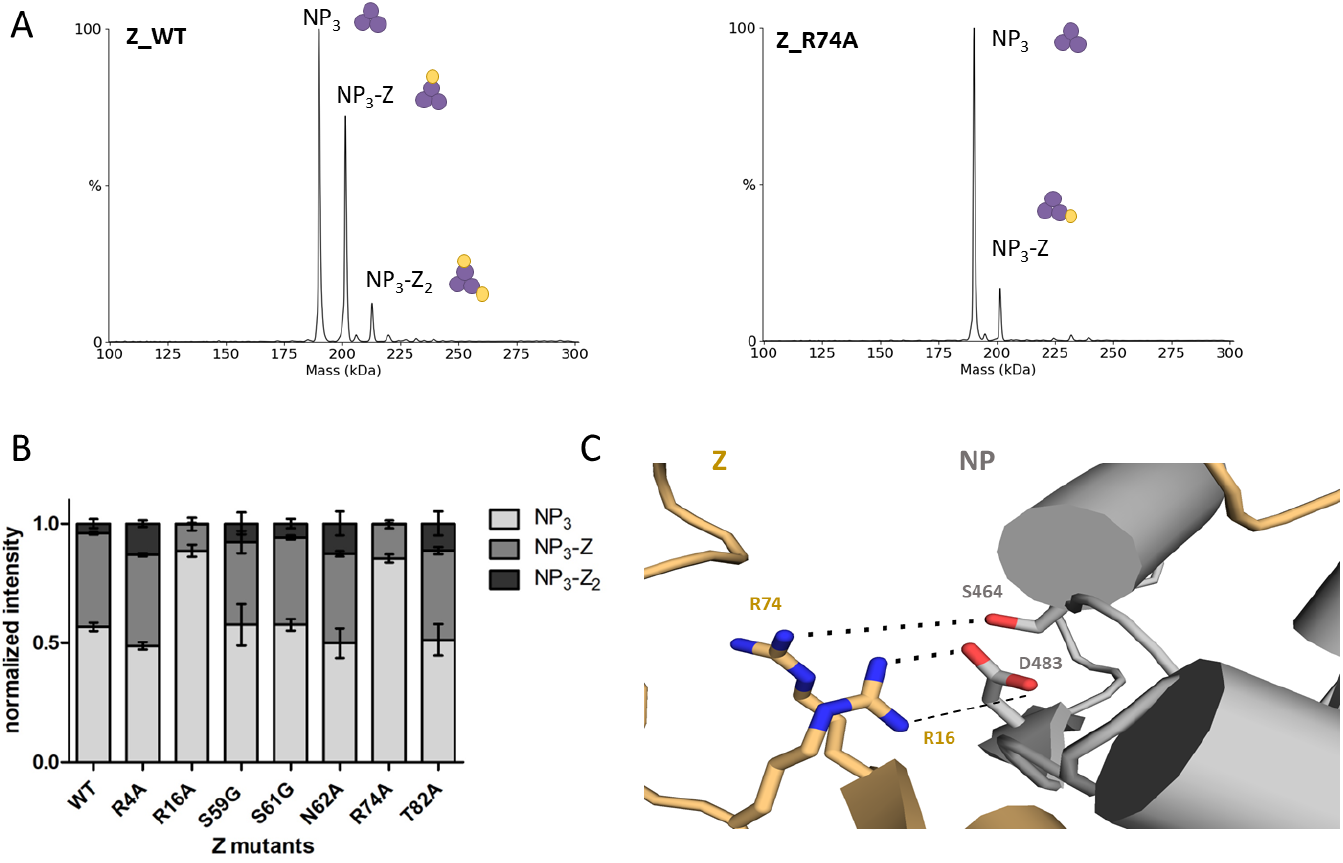
Z mutant screening identifies important residues for NP-Z interaction. The different Z mutants were recombinantly expressed and purified. (A) nMS screening of the Z mutants together with NP (6 μM) in a 1:3 (NP:Z) molar ratio in a 150 mM ammonium acetate solution at pH 7.5. (B) The sum of the intensities of every mass species of one measurement was normalized to 1. The normalized intensities for every measurement were plotted. Error bars represent standard deviation of at least 3 independent measurements. Measurements with the mutants R16A and R74A show reduced abundance of NP-Z complex species compared to wildtype (WT). (C) Detailed closeup to the Z residues R74 and R16 in the AlphaFold Multimer model. Z residues R74 and R16 are in close proximity to NP-S464 and NP-D483, respectively.

Taken together, our structural integrative approach including AI complex prediction, HDX-MS and mutational analysis in combination with nMS lead to detailed information about the direct NP-Z interaction. Z seems to bind at the C terminus of NP, where arginine R16 and R74 of Z are likely important for the NP-Z interaction.

### RNA triggers disassembly of trimeric NP

To investigate the effect of RNA binding on the NP quaternary structure, NP_3_ was incubated with short RNAs. As no sequence specificity of NP for RNA has been described so far, we used RNAs of different sequences and lengths. First, we mixed a 9 nt RNA (Tab. S2) in an equimolar ratio with NP and monitored the changes in quaternary structure over at least 500 s in the electrospray ionization (ESI) capillary (Fig. 6A). The acquisition started approximately 30 s after mixing the components and mounting the capillary. During the measurement, the fractions belonging to the NP_3_ decreased whereas peaks assigned to a mass of 66.08 +/-0.02 kDa, corresponding to NP monomer bound to one RNA molecule (NP_1_-RNA_1_), increased. Importantly, we also observed NP_3_ interacting with one RNA molecule (193.00 +/-0.09 kDa) at intermediate timeoints (Fig. 6B). As the RNA binding groove of NP is inaccessible in the trimeric ring conformation the observed complex of one NP trimer and one RNA molecule (NP_3_-RNA_1_) presumably constitutes an open conformation of NP_3_, allowing the RNA to bind. After 500 s the NP_3_ was mostly disassembled. To further support our results, we used negative-stain electron microscopy to confirm the disassembly of NP_3_ in the presence of RNA. As in our nMS data, we observed NP mainly as a ring-shaped trimer in the absence of RNA (Fig. 6C, S7A). In the presence of RNA, however, the morphology changes to smaller rounded particles and more heterogenous species as NP_3_ is not detectable anymore (Fig. 5C, S7B). Furthermore, we validated our results by mass photometry. We observed NP monomer formation in presence of RNA (Fig. S8). We conclude that RNA is sufficient to trigger conformational changes leading to the disassembly of the trimeric ring structure of NP into RNA-bound NP monomers.

**Figure 6:**
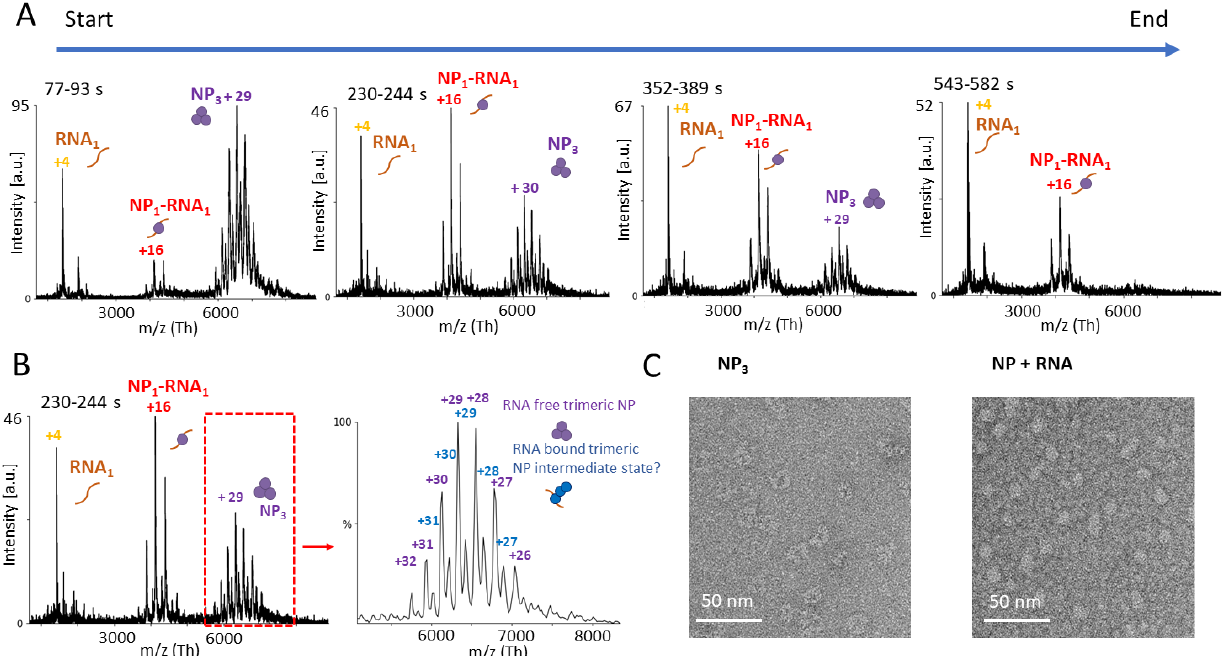
RNA triggers the dissociation of NP_3_ into monomers. NP_3_ and a 9 nt RNA were mixed together in a 150 mM ammonium acetate solution at pH 7.5 on ice in a 1:1 (NP_1_:RNA) molar ratio. The mixture was placed into an ESI capillary and the measurement was started approximately 30 s after mixing RNA and NP together. Spectra were recorded over 550 s. (A) Each diagram represents combined spectra recorded at 77-93 s, 230-244 s, 355-389 s, and 543-582 s with the intensity in arbitrary units (a.u.). Main charge states are labeled. Shown is one representative measurement of at least 3 independent measurements. (B) Recorded spectrum at timepoint 230-244 s. Zoom into the *m/z* range between 5000 and 8000 shows an RNA bound NP_3_ (193.00 +/-0.08 kDa) species. (C) NP at 160 nM was mixed with a single-stranded 25 nt RNA at a 1:2 molar ratio (NP:RNA) (Tab. S3) and incubated at room temperature for approximately 15 min. NP was applied to glow-discharged carbon-coated copper grids and stained with uranyl acetate immediately before imaging. Images were collected with a transmission electron microscope.

Next, we used a slightly longer RNA of 12 nts (Tab. S3) to investigate a potential assembly of RNA bound NP monomers, corresponding to the beginning of higher order NP-RNA assemblies. We again mixed the components and determined the complex species over a duration of 45 min in an ESI capillary. The measurement was started approximately 30 sec after mixing RNA and NP together. We first tested an equimolar ratio of NP to the RNA of 12 nts (Fig. 7). After the start of the measurement 3 species were visible: (i) NP_3_, (ii) a trimeric NP bound to one RNA molecule (NP_3_-RNA_1_,193.63 +/-0.09 kDa) and (iii) unbound RNA (3.9 kDa) (Fig. 7B). One minute later, a signal corresponding to an NP monomer bound to one RNA molecule (67.03 +/-0.05 kDa) became visible. Between 5 and 30 min after the start of the measurement NP-RNA complexes assembled corresponding to NP_2_-RNA_2_ (134.03 +/-0.05 kDa), and NP_3_-RNA_3_ (201.17 +/-0.13 kDa) species. As in the previous experiment, the signal for the ring-structured NP_3_ decreased over time. The NP_3_-RNA_3_ complexes were most abundant after around 30 min of measurement.

**Figure 7:**
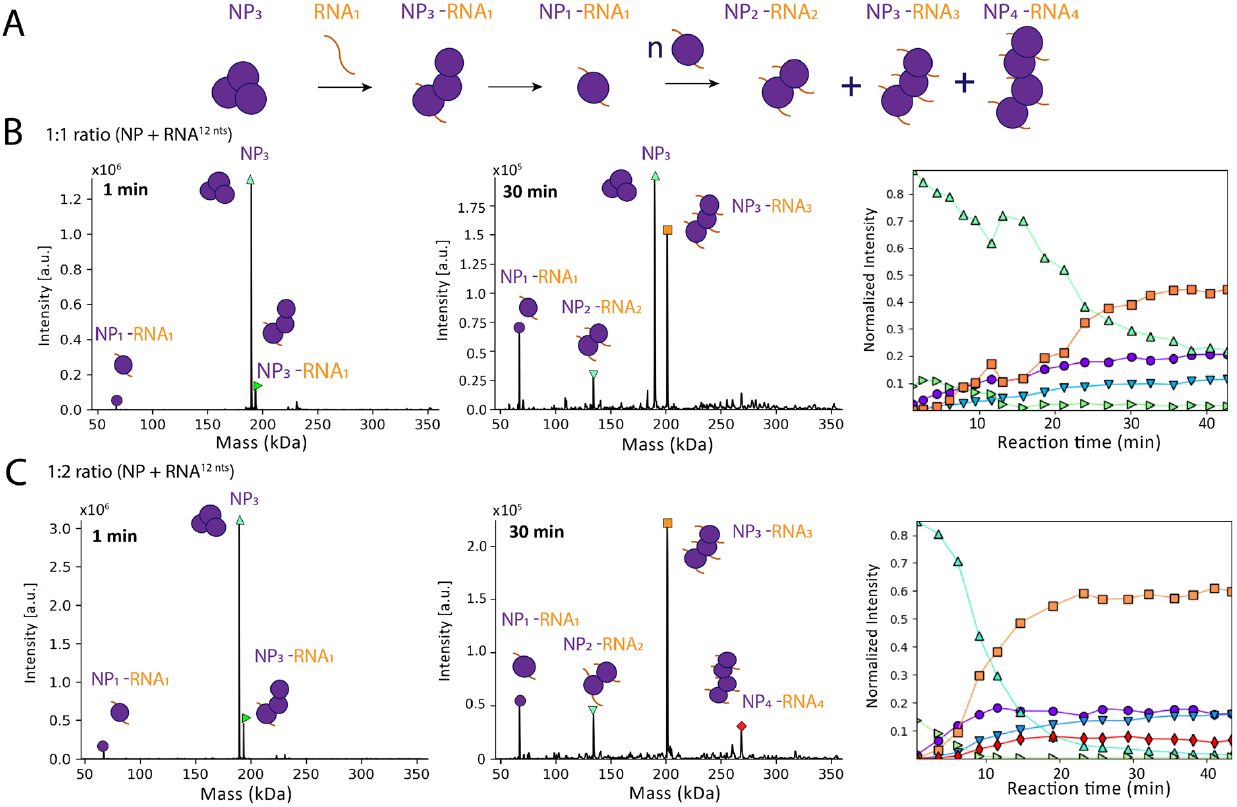
NP_3_ dissociation and step-wise assembly of nucleocapsids. (A) Schematic summary of NP_3_ dissociation and NP-RNA complex assembly over time as detected in the experiment. (B) Representative measurement of NP (9 uM) together with a 12 nt RNA in a 1:1 (NP:RNA) molar ratio recorded for 45 min in a 150 mM ammonium acetate solution at pH 7.5. Spectra of all timepoints were deconvoluted with Unidec [30] shown are spectra at 1 min after starting the measurement (left) and at 30 min after starting the measurement (middle) with the intensity in arbitrary units (a.u.). The cartoons represent the complexes of NP and NP-RNA. Masses corresponding to NP_3_ (189.56 +/-0.02 kDa), NP_3_-RNA_1_ (193.63 +/-0.09 kDa), NP_1_RNA_1_ (67.03 +/-0.05 kDa), NP_2_-RNA_2_ (134.03 +/-0.05 kDa), and NP_3_-RNA_3_ (201.17 +/-0.13 kDa). The normalized intensity for every mass species from the deconvoluted spectra were plotted according to the different timepoints. The sum of all mass species of one timepoint was normalized to 1 (right). (C) Representative measurement of NP together with a 12 nt RNA in a 1:2 (NP:RNA) molar ratio. Spectra were analyzed as in (B). Masses corresponding to NP_3_ (189.56 +/-0.02 kDa), NP_3_-RNA_1_ (193.63 +/-0.09 kDa), NP_1_-RNA_1_ (67.03 +/-0.05 kDa), NP_2_-RNA_2_ (134.03 +/-0.05 kDa), NP_3_-RNA_3_ (201.17 +/-0.13 kDa) and NP_4_-RNA_4_ (268.25 +/-0.05 kDa).

Notably, NP:RNA ratios of the complexes were always stoichiometric after initial disassembly of the ring-structured NP_3_ via NP_3_-RNA_1_. We increased the RNA concentration to a molar ratio of 1:2 (NP:RNA) to further evaluate the formation of higher-order RNA-NP complexes, which was significantly faster compared to using an equimolar ratio (Fig. 7C). Although the NP_3_ and the NP_3_-RNA_1_ complex (193.3 +/-0.09 kDa), likely an intermediate, were observable at early timepoints of the measurement, the decay rate of the trimeric ring structure was higher than observed in the previous experiment with an equimolar NP:RNA ratio and NP_3_ had completely disappeared after 40 min. In this experiment, we observed formation of NP_1_RNA_1_ (67.03 +/-0.05 kDa), NP_2_-RNA_2_ (134.03 +/-0.05 kDa), NP_3_-RNA_3_ (201.17 +/-0.13 kDa) and NP_4_-RNA_4_ (268.25 +/-0.05 kDa) complexes. Small fractions of NP_5_-RNA_5_ (335.78 kDa, FWHM: 0.90 kDa), and NP_6_-RNA_6_ (393.18 kDa, FWHM: 1.12 kDa) and) were also detectable (Fig. S9A). In addition to the 12 nt RNA-NP complexes, we detected NP monomers bound to truncated RNAs (RNA_tr_). As we did not observe any RNA_tr_ as free species, the NP_1_-RNA_tr_ complexes were most likely a result of an onset of CID, clipping off nucleotides not covered by the RNA-binding groove of NP (Fig. S9B). These NP_1_-RNA_tr_ complexes corresponding to 8-9 bases bound to NP were observed with different RNA substrates of varying lengths. This is in line with previously reported crystal structures of NP bound to RNA, where 6 nts were fit into the RNA-binding groove [9].

Furthermore, we can rule out that the assembly between NP molecules happened via base pairing of overhanging RNA, as with our acceleration voltage settings only single RNAs were observed (Fig. S9).

Altogether, these results show the higher-order assembly of NP and RNA possibly resembling RNP formation during viral replication. Based on our data, we hypothesize that, as a first step, one molecule of NP within the RNA-free NP trimer binds to RNA leading to an opening of the ring-like conformation, which is less stable than the closed ring-like trimer. After this trigger, the open trimer further disassembles into RNA-bound NP monomers. Starting from that, higher-order NP-RNA complexes are then formed (Fig. 7A). We could demonstrate that this process is dependent on the RNA concentration as an excess of RNA relative to NP led to a faster trimer disassembly and higher-order RNP-like NP-RNA complex formation (Fig. 7B).

### NP can bind Z and RNA simultaneously

As described above, Z binds to the C-terminal domain of NP whereas the RNA binding pocket of NP is located in the N-terminal domain. Co-immunoprecipitation data has previously shown for other arenaviruses that NP and Z, either directly or mediated by other factors, interact with each other [6, 14, 31]. It has been hypothesized that the NP-Z interaction mediates virion assembly by recruiting viral RNPs to the plasma membrane. We therefore investigated if LASV NP could indeed bind to Z and RNA simultaneously. Mixing a 9 nt RNA, NP and Z in a 1:1:2 ratio for subsequent nMS measurements, the NP_3_ ring conformation again dissociated into RNA-bound monomers (Fig. 8A) (66.08 +/-0.03 kDa), some of which were associated with Z (77.38 +/-0.06 kDa) (Fig. 8B). CID-MS/MS confirmed these results (Fig. 8CD). In both cases the RNA remained bound to NP at high collision voltage whereas Z dissociated. This indicates a strong ionic interaction between RNA and NP compared to a weaker interaction between NP and Z. This observation fits with our proposed model of NP-Z interaction, which shows a rather small protein-protein interface between the C-terminal domain of NP and Z (Fig. 4B). The affinity between wild-type NP and Z for the NP_1_-Z_1_-RNA_1_ complex appeared to be in the same range when using the NP-R52A trimerization mutant with a K_D_ around 30 μM. Taken together, these results show that the NP_1_-RNA_1_ complex can indeed interact with Z, supporting the idea that Z mediates the recruitment of RNPs to the plasma membrane by direct interaction of Z with RNA-bound NP.

**Figure 8:**
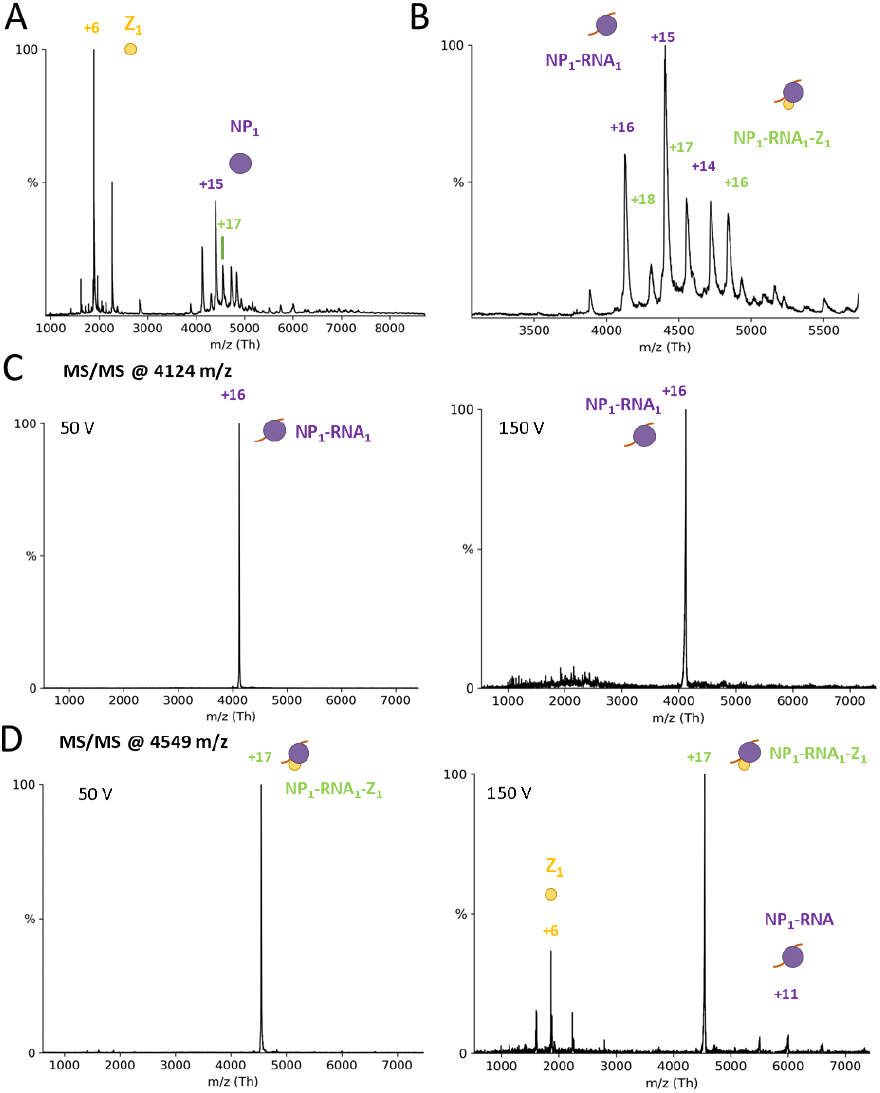
NP can bind Z and RNA simultaneously. (A) Full nMS spectrum of NP with Z and 9 nt RNA in a 150 mM ammonium acetate solution at pH 7.5. NP_3_ completely dissociates into RNA-bound monomers. (B) Closeup of the *m/z* range 3,000-6,000 for determination of masses corresponding to NP-Z and RNA-NP-Z complexes. MS/MS of peaks corresponding to NP-Z isolated at 4,124 *m/z* (C) and NP_1_-Z_1_-RNA_1_ isolated at 4,549 *m/z* (D) at low and high acceleration voltage in the collision cell. CID products for the NP_1_-Z_1_-RNA_1_ complexes led to masses corresponding to Z monomers and NP_1_-RNA_1_ complexes, whereas no CID products for the NP-RNA complex were observed at higher acceleration voltage.

### NP-Z interaction is pH-dependent

If the NP-Z interaction is responsible for RNP recruitment into budding virions, we wondered what triggers the release of the RNPs upon virus entry and membrane fusion. A potentially important factor could be the pH as LASV is taken up via the endosomal pathway. It was recently shown that acidification of Lassa virions occurs before membrane fusion via reorganization of the glycoprotein complex [32, 33]. A low pH could lead to dissociation of RNPs from the Z protein lining the inner side of the viral membrane. Therefore, we evaluated the pH-dependence of the NP-Z interaction by incubating the proteins at varying pH between 5 and 7.5 and subsequently measured NP-Z complex formation by nMS (Fig. 9A). The first step was to evaluate the protein homogeneity after changing the buffer to lower pH. NP showed a similar oligomerization behavior at pH 5 and pH 7.5, with the masses still corresponding to a trimer (Fig. 9B). In contrast, Z started to homo-oligomerize after changing the pH to 5 forming up to homohexameric complexes (Fig. 9B). This is in line with an observation on influenza matrix protein that also undergoes oligomerization at low pH [34]. However, we observed very similar protein concentrations at both pH 5 and pH 7.5 after buffer exchange and high-speed centrifugation indicating that no unspecific protein aggregation happens during buffer exchange. To test the pH-dependence of the NP-Z interaction, we recorded 3 independent spectra of a mixture of NP and Z at a 1:3 (NP:Z) molar ratio at pH 5, 5.5, 6.5 and 7.5. We found that almost no NP-Z interaction was observed at pH 5 while complex formation at Ph 7.5 was as observed in previous experiments (Fig. 9C). At pH values between 5.5 and 6.5 NP-Z interaction was also less abundant compared to pH 7.5 (Fig. 9D). These experiments show that the interaction between NP and Z is highly pH-dependent. This supports our hypothesis that the endosomal acidic pH reduces NP-Z interaction and, as a result, RNPs are released from the viral matrix.

**Figure 9:**
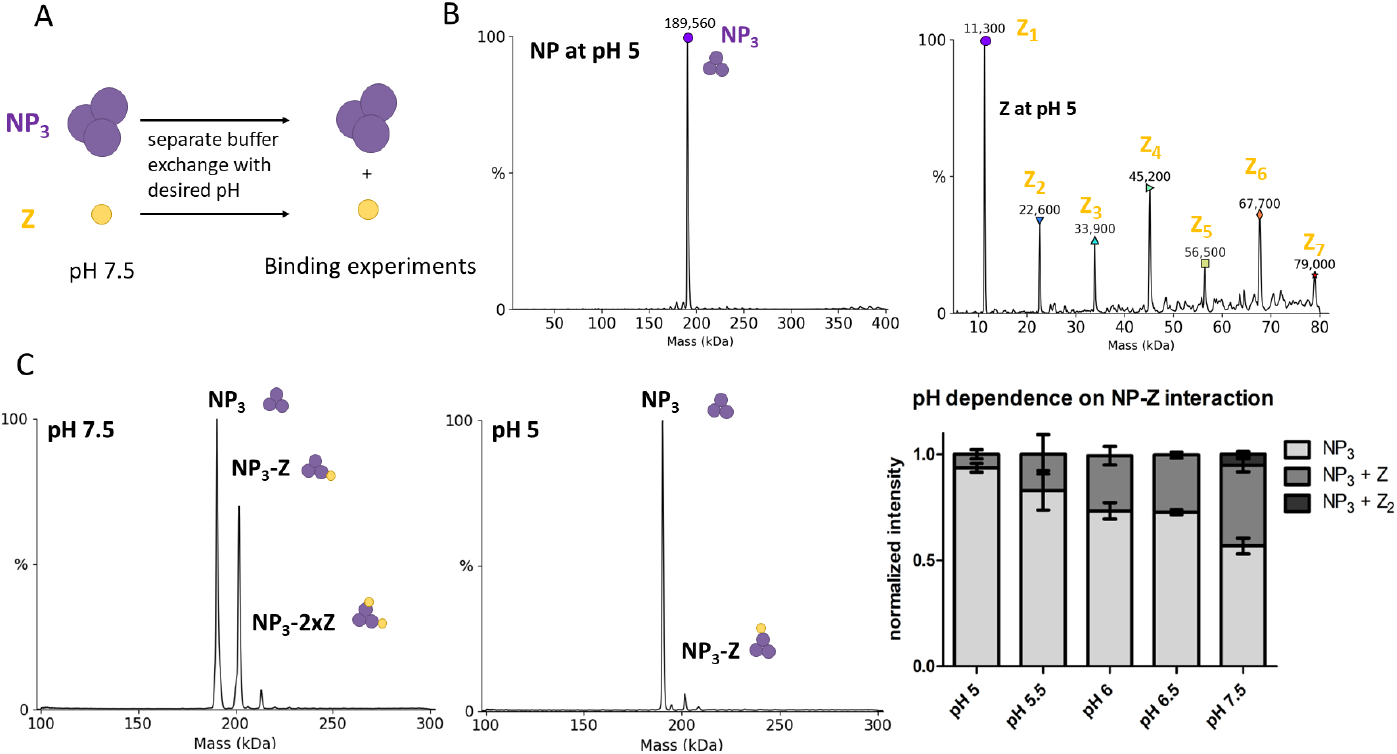
NP-Z interaction is less efficient at lower pH. (A) Scheme of the pH-dependent interaction assay. Proteins were separately exchanged to 150 mM ammonium acetate surrogate with adjusted pH. NP (6 μM) and Z with the same pH were incubated in a 1:3 (NP:Z) molar ratio and measured by nMS. (B) Deconvoluted spectra of NP and Z after changing the buffer to pH 5. NP oligomerization status was unaltered whereas Z starts to oligomerize at pH 5 and 5.5. (C) Spectra of NP and Z in a 1:3 molar ratio at different pH. (D) The normalized intensity for every mass species from the deconvoluted spectra were plotted according to the different pH conditions. The sum of all mass species of one measurement was set to 1. Error bars representing standard deviation of at least 3 independent measurements.

**Figure 10:**
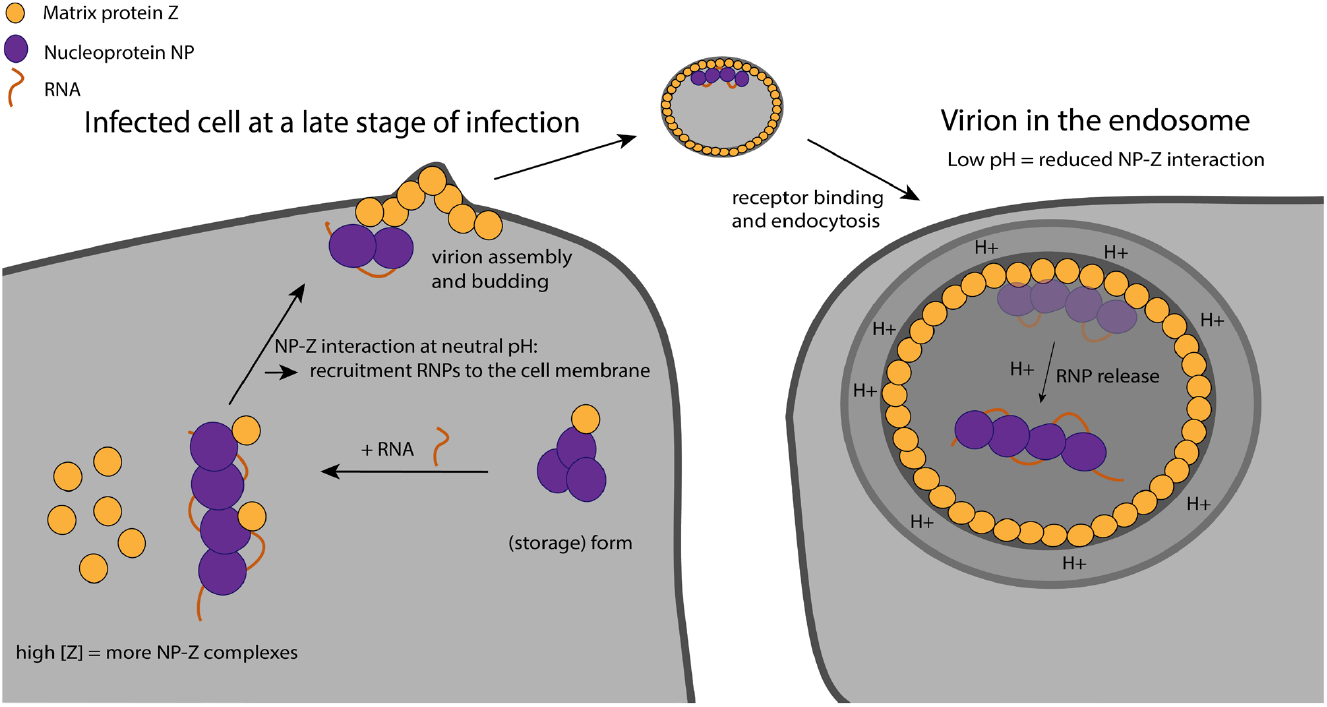
Proposed model of the NP-Z interaction during the viral life cycle. At a late stage of infection, the cellular concentration of Z is high favoring NP-Z interaction. This leads to recruitment of the RNPs to the cell membrane by Z for virion assembly (left). During virus entry into the host cell via the endosomal pathway, the lower pH leads to dissociation of NP from Z, which coats the inner virion membrane. This leads to the release of the RNPs into the cytosol.

## Discussion

NP is known to form trimers in the absence of RNA and during recombinant purification with its atomic structure solved [8]. The trimeric NP can be isolated from RNP containing cells alongside full RNP complexes demonstrating existence of this complex in infected cells [23, 35]. It was speculated that the trimer serves as a storage form for NP in cells. The ability of NP to trimerize was found to be essential for viral genome replication and transcription but was dispensable for interferon antagonism and NP recruitment into budding virions [23]. It was assumed that the NP trimer is a functional equivalent of the NP-phosphoprotein complex of non-segmented negative-strand RNA viruses preventing unspecific RNA binding to the NP [9, 36]. Structurally, the trimeric ring conformation of arenavirus NP is unable to bind RNA. To enable this, the C-terminal domain must first rotate away from the RNA-binding pocket allowing one helix to become partly unstructured (α5) and a second helix (α6) to shift away from the RNA binding pocket [9, 37]. To induce this conformational change, it was speculated that specific (host-) factors are needed [9]. In our experiments, we demonstrated that short RNAs are sufficient to induce this conformational change and trigger NP trimer dissociation. Yet, how does NP primarily associate with viral RNA instead of host RNA? We did not observe any sequence specificity as NP bound to all random RNAs used in our experiments, which is in line with other groups reporting randomly bound RNAs to LASV NP [9, 25]. However, it cannot be ruled out that the LASV RNA genome contains some specific structural and sequence motifs (packaging signals) or additional modifications that could lead to a lower threshold for RNA binding. These signals would have to be different in the genomic RNA compared to viral mRNA, which is usually not encapsidated. It also remains unclear whether and how specific recruitment of all genome segments is controlled in bunyaviruses as specific RNA secondary structures, as present in influenza viruses [38], seem to be missing. For Rift Valley fever virus, a non-selective recruitment of viral genome segments into budding virions was found to be the most likely scenario [39-41]. Another factor favoring viral RNA binding could be the local RNA concentration. In our experiments, we used relatively high RNA concentrations (1:1 or 1:2 molar ratio) and observed the dissociation and assembly process in real time, pointing to rather slow RNA binding kinetics. Several scenarios are conceivable in support of the idea of local viral RNA concentration as a key determinant for preferred and efficient packaging of viral genomic RNA:

(i) Direct interaction of NP with the L protein for close proximity to nascent viral RNA. NP is a critical component of the arenavirus replication machinery. A direct interaction of NP and L was suggested based on co-immunoprecipitation studies, although an interaction mediated by RNA or host factors could not be ruled out in this case [42, 43].

(ii) Indirect interaction of NP with L during viral genome replication via a host protein. This has been observed for influenza virus polymerase complex, where the host acidic nuclear protein (ANP32) enables L-L interaction and also binds to NP to support encapsidation of nascent viral RNA and re-encapsidation of the template [44, 45].

(iii) Formation of local microenvironments for viral genome replication and transcription within the cell. Bunyavirus NPs have been detected in non-membranous compartments such as processing bodies and stress granules in cells [46-49]. Other studies reported NP containing cytosolic puncta forming upon arenavirus infection associated with cellular membranes. These puncta were concluded to contain full-length genomic and antigenomic RNAs along with proteins involved in cellular mRNA metabolism reminiscent of classical replication-transcription complexes [50]. Interestingly, phosphorylation of LCMV NP at position T206 was found to be critical for the formation of these cytosolic puncta [51]. However, there is no evidence that the L protein is indeed present in these puncta and the presence of these local puncta would not provide an explanation on why mRNAs are not encapsidated. It therefore remains unclear what exactly determines viral RNA packaging by NP in addition to the presence of the RNA molecule itself.

In our experiments, we could observe NP trimer dissociation into RNA-bound NP monomers, which then assembled to higher-order NP-RNA structures. NP-RNA higher-order complex formation seemed to require a certain RNA length, indicating an additional stabilizing effect of the RNA or even a secondary RNA binding site in NP as observed for other segmented negative-strand RNA viruses [52]. Interestingly, we observed a complex of one RNA molecule bound to the NP trimer (NP_3_-RNA_1_) although according to our interpretation of the available structural data, RNA binding is not possible in the trimeric ring conformation [8]. We propose that the observed NP_3_-RNA_1_ state is therefore likely a disassembly intermediate. Mechanistically, we hypothesize that the trimer needs to open up first, allowing for one RNA-binding site to be occupied, while the NP molecules are still connected to each other. The open conformation is less stable and therefore the trimer quickly disassembles into monomers which immediately bind RNA. This indicates overall a cooperative dissociation mechanism.

It has long been known that arenavirus NP and Z are acting together, based on cellular experiments, such as VLP assays, co-immunoprecipitation and colocalization studies [10, 12, 14, 53]. Here we showed that NP and Z directly and independent of any host factor interact with each other. Importantly, monomeric NP in presence or absence of RNA has a substantially higher affinity for Z compared to the NP_3_ storage form, which is not recruited to the membrane. We mapped the NP-Z interaction site with an integrative structural approach of nMS, HDX-MS and Alphafold Multimer-powered complex prediction. Our data point towards Z binding to the C-terminal domain of NP likely involving the side chains of R16 and R74 of Z. Previous co-immunoprecipitation studies already suggested the C-terminal domain of NP to be important for the interaction with Z protein, either directly or mediated by cellular factors [6]. Several conserved Z residues, including R16 and T73, were also demonstrated to be important for inhibition of viral RNA synthesis and virus-like particle (VLP) infectivity although R16 and T73 are not involved in the binding of Z to the L protein [20, 54]. This points towards the importance of R16 and T73 conservation for Z localization and/or proper folding. However, we could not detect a difference in the protein folding comparing Z R16A to the wild-type Z *in vitro*.

It was hypothesized that Z residues R16, T73 and others were involved in RNP recruitment via binding to NP and/or viral RNA [54]. Our experimental data and the proposed model support this hypothesis as we find R16 and T73 to be located in the interface between NP and Z, while we did not detect an interaction between Z and RNA *in vitro* as observed for the matrix protein of other negative strand RNA viruses [55]. Upon RNA packaging, RNPs are recruited into nascent virions and Z has been shown to be critical for budding of most arenaviruses [56]. While Z alone is sufficient to drive the formation of VLPs, the process is significantly enhanced when NP is present [14]. For Tacaribe virus, the so-called late-domain motifs ASAP and YxxL of Z have an influence on the NP incorporation, with the Y56 in the YxxL motif being critical. These data suggest a role of the late-domain motif YxxL in the NP-Z interaction [57]. Interestingly, although the YxxL motif is also present in the LASV Z, the Y48 of the YxxL motif is not in the proposed NP-Z interface in our model but rather on the opposite side of the Z molecule. However, the proposed location of this motif in our model is in agreement with another study reporting the binding of Z to a host protein required for budding via the YxxL motif [58].

The data shown here support the concept, that the NP-Z interaction is important for the recruitment of RNPs to the plasma membrane and incorporation into nascent virions. We demonstrate that RNA-bound NP is able to bind Z as would be expected for RNP recruitment.

Arenavirus cell entry is mediated by the glycoprotein complex. After receptor binding, virions enter the host cell via the endosomal pathway [59, 60]. Our data show that the interaction between NP and Z is highly pH-dependent with a decreased NP-Z interaction at pH values lower than pH 7.5. Therefore, it is conceivable that RNP-Z dissociation triggered by a lower pH in the endosome leads to the release of RNPs into the cytosol after viral membrane fusion. Unlike in influenza viruses, arenaviruses do not have proton channels in their lipid envelope that actively acidify the virion upon endocytosis. However, it has recently been shown for LASV that protons passively cross the viral membrane which leads to acidification of virus interior prior to membrane fusion [32, 33]. Similar observations have been reported for Ebola virus, were acidification of virions leads to viral matrix reorganization [61]. Besides the pH-dependent NP-Z interaction, we observed that Z starts to oligomerize between pH 5 and 5.5. We hypothesize, that stronger Z-Z interaction at low pH, as indicated by oligomerization, further reduces availability of Z for the low affinity Z-NP interaction. This is in line with several studies that report pH as a critical factor for matrix protein oligomerization and protein conformation [34, 62-65]. Matching our observations, a lower pH at around 5.4 also induced weakening of influenza virus matrix protein M1-vRNP interactions [66].

Our findings support a model, in which viral RNA is necessary and sufficient to transform trimeric NP into monomeric RNA-bound, RNP-like NP assemblies. Z then binds to the RNP at neutral pH in the cytosol or a cellular microenvironment via direct interaction with NP. This interaction facilitates targeted transport of the RNP to the plasma membrane at high cellular concentration of Z protein during late stages of the viral “life” cycle and subsequent virion budding. Upon virus entry into a host cell, the RNPs are released into the cytosol from the virion matrix triggered by the low pH in the endosome and following membrane fusion. Subsequently viral genome replication and transcription take place.

## Materials and Methods

### Expression and purification

Lassa Z (AV strain) was expressed with a pOPIN-J vector with an N-terminal GST tag and purified as described in [16]. Mutations (if indicated) were introduced by mutagenic PCR before cloning into the pOPIN-J Vector and expressed and purified according to the WT. Lassa NP (AV strain) was expressed with a pOPIN-M vector with an N-terminal maltose binding protein (MBP) fusion protein. Mutations (if indicated) were introduced by mutagenic PCR before cloning into the pOPIN-M. Expression was performed in BL21 in *E. coli* strain BL21 Gold (DE3) (Novagen) at 37°C and grown in 1-liter cultures in terrific growth media to an optical density at 600 nm of 1.0. Cells were induced with 250 μM isopropyl-β-d-thiogalactopyranoside and incubated overnight at 17°C. Cells were harvested by centrifugation and lysed by sonification in 50 mM NaH_2_PO_4_, pH 8.0, 500 mM NaCl, and 5% (w/w) glycerol, 1 mM phenylmethylsulfonyl fluoride, 0.05 % (v/v) TritonX-100, 0.025% (w/v) lysozyme and 2 mM dithiothreitol. The lysate was clarified by centrifugation at 30,000 *g* for 30 min at 4°C and incubated for 45 min with amylose beads (GE Healthcare). The loaded protein was cleaved on column by GST-tagged 3C protease at 4°C overnight. The NP without MBP containing buffer was diluted to reach 150 mM NaCl concentration by adding 20 mM NaH_2_PO_4_ pH 8 and loaded on a heparin column (GE Healthcare) to remove bound bacterial RNA. NP was eluted by an NaCl gradient and concentrated for further purification by size-exclusion chromatography on an SD200 column (GE Healthcare) in 50 mM Tris(HCl) pH 7.5, 150 mM NaCl, 5% (w/w) glycerol, 2 mM dithiothreitol. Proteins were concentrated using filter units (Amicon Ultra, 30,000 Da cutoff), flash frozen in liquid nitrogen and stored at -80°C.

### Native MS sample preparation

Proteins for native MS analysis were buffer-exchanged into 150 mM ammonium acetate (99.99 % purity) at pH 7.5 (if not indicated otherwise) by at least three cycles of centrifugation at 13,000 *g* and 4°C in centrifugal filter units (Amicon 3,000 or 30,000 molecular cutoff and 500 μl volume). Proteins were centrifuged at 20,000 *g* for 15 min at 4 °C to clear the sample from possible protein aggregates occurring during the buffer exchange. Protein concentration after buffer exchange ranged from 5 to 100 μM. Proteins were mixed with or without ligands on ice and subsequently introduced to a mass spectrometer.

Samples were introduced to the mass spectrometer by gold-coated capillaries made from borosilicate glass tubes (inner diameter: 0.68 mm, outer diameter: 1.2 mm; World Precision Instruments). Capillaries were pulled in a two-step program using a micropipette puller (P-1000, Sutter Instruments) with a squared box filament (2.5 mm by 2.5 mm, Sutter Instruments) Capillaries were afterwards gold-coated using a sputter coater (5.0 × 10^−2^ mbar, 30.0 mA, 100 s, three runs to vacuum limit 3.0 × 10^−2^ mbar argon, CCU-010, Safematic)

### Native MS data acquisition

Data acquisition was performed on a nanoESI Q-ToF II mass spectrometer (Waters/Micromass) modified for high mass experiments in positive ion mode [67]. Samples were ionized with voltages at the capillary of 1450 V and at the cone of 150 V. Pressure of the in-source region was constantly at 10 mbar. Pressure in the collision cell was at 1.5 × 10^−2^ mbar argon. Quaternary structure of protein complexes was determined in MS/MS mode with acceleration voltage ranging between 50-400 V across the collision cell. The quadrupole profile was 1,000 to 10,000 *m/z* throughout all experiments. Cesium iodide (25mg/ml) in aqueous solution was used to calibrate the instrument and the data.

The NP-RNA assembly experiment (Fig. 7) was performed on a Q Exactive Ultra-High Mass Range (UHMR) Orbitrap mass spectrometer (Thermo Fisher Scientific) in positive ion mode with a mass range from 350 to 20.000 *m/z* and a nano ESI source. Following parameters were used: Capillary voltage was kept at 1.2 kV, source temperature at 50 °C, noise level parameter at 4.64, S-lens radio frequency level 200, in source CID at 5 eV, resolution at 6250 setting value, ion injection time of 100 ms. Nitrogen was used in the HCD cell with a trapping gas pressure of 5 setting value with ultra-high vacuum readout at around 2.5 × 10^−10^ mbar. HCD energy setting was at 100. Instrument detector parameters were set to low *m/z*. 10 microscans were summed into 1 scan.

### Data analysis for native MS

Raw data was analyzed with MassLynx V4.1, (Waters) for QToF2 datasets or Xcalibur for UHMR datasets by combining spectra and exporting them into txt file format. Following evaluation regarding full width at half maximum (FWHM) was done with mMass (Martin Strohalm [68]). Experimental mass, area under the curve (AUC) and visualization was evaluated with Unidec (Michael T. Marty [30]). K_d_ determination was calculated after [69]. The values for all recorded masses and FWHM resulted from at least 3 independent measurements and are provided in Table S4.

### HDX-MS

NP protein (50 pmol) was mixed with the Z protein in a 1:5 molar ratio (NP:Z). The deuterium exchange reaction was started by a 1:9 dilution into 99% deuterated buffer (40 mM Tris(HCl) pH 7.5, 150 mM NaCl) at 25°C. The exchange reaction was quenched after the following time points: 15 s, 1 min, 10 min, 1 h, 6 h. The quenching was initiated by a 1:1 addition of ice-cold quench buffer (1 M glycin pH 2.3). After that, samples were flash-frozen in liquid nitrogen. Additionally a fully deuterated (FD) control was prepared were NP was diluted 1:9 in 99 % deuterated 40 mM Tris(HCl) pH 7.5, 150 mM NaCl, 6 M urea buffer. The reaction was quenched after 24 h incubation at room temperature as described above. All timepoints were analyzed in three technical replicates.

The samples were shortly centrifuged after thawing and injected onto a cooled (2°C) HPLC System (Agilent Infinity 1260, Agilent Technologies, Santa Clara, CA, USA) which includes a home packed pepsin column (IDEX guard column with 60 μl Porozyme immobilized pepsin beads, Thermo Scientific, Waltham, MA, USA), in a column oven 5°C) a peptide trap column (OPTI-TRAP for peptides, Optimize Technologies, Oregon City, OR, USA) and a reversed-phase analytical column (PLRP-S for Biomolecules, Agilent Technologies, Santa Clara, CA, USA). Peptide digestion was carried out online at a flow rate of 75 μl/min (0.4 % formic acid in water) and washed after every run with 2 M urea, 2 % acetonitrile, 0.4 % formic acid. Peptide separation prior MS analysis was performed on an analytical column at a flow rate with 150 μl/min and a gradient of 8-40 % buffer B in 7 min (buffer A: 0.4% formic acid in water, buffer B: 0.4 % formic acid in in acetonitrile). The analytical column was washed with 100 % of buffer B after every sample. The HPLC was connected to an Orbitrap Fusion Tribrid in positive ESI MS only mode (Orbitrap resolution 120,000, 4 microscans, Thermo Scientific, Waltham, MA, USA).

Peptide identification was evaluated with 100 pmol of non-deuterated samples. A gradient from 8-40 % buffer B in 27 min was used for the analytical column with a flow rate of 150 μl/min. MS analysis was perfomed in data-dependent MS/MS acquisition mode (Orbitrap resolution 120,000, 1 microscan, HCD 30 with dynamic exclusion). Raw data is available in the pride database entry.

### HDX Data analysis

Peptides were searched against the protein sequence in MaxQuant (version 2.1.2.0) with the Andromeda search engine[70]. Following MaxQuant settings were used: digestion mode was set to unspecific and peptides between 5 and 30 amino acids length were accepted. The minimum score for identification of peptides was 20. The default mass tolerance for precursor (4.5 ppm) and fragment (20 ppm) ions were used according to the Thermo Orbitrap instrument. The retention time for peptides of the HDX runs were adjusted to the shorter gradient of the analytical column compared to the longer gradient of the peptide identification runs.

Datasets were analyzed according to deuterium uptake of peptides via the automated centroid analysis by HDExaminer Version 3.3 (Sierra Analytics). All peptides were checked manually based on the presence of overlapping peptide envelopes, correct retention time, *m/z* range, and charge state. All peptides at different states and timepoints are plotted as wood plot (Fig. S5) and difference heatmap according to the atomic range which considers partly overlapping peptides with no difference (Fig. S6). A Summary of all experimental conditions and statistics can be found in Table S2.

GraphPAD Prism (GarphPAD Software, Inc) and PyMOL (Schrödinger) software were used for visualization. All evaluated data points are available in the pride database entry.

### Negative-stain electron microscopy

Purified NP at 0.01 mg/mL was applied to glow-discharged carbon-coated copper grids (Electron Microscopy Sciences) and stained with uranyl acetate immediately before imaging. Transmission electron microscopy images were collected using a Talos L120c microscope (120 kV) with a LaB_6_ thermionic source at x92,000 magnification and a defocus of between -0.2 and 0.5 μm. For the NP + RNA condition, NP at 0.01 mg/mL was mixed with a single-stranded 25mer (Table S3) at a 2 molar-excess to the NP monomer. The NP + RNA was incubated at room temperature for approximately 15 minutes and then imaged as for the NP without RNA condition.

### Structural modelling of LASV NP-Z complex

To model the NP-Z complex, we used the ‘custom’ mode of AlphaPulldown [71], a python package built upon AlphaFold [27] and AlphaFold Multimer [28]. Both the initial release of AlphaFold Multimer (version 2.1.0) and the latest version (version 2.3.0) were used. All parameters were set to default except for the max_recycles, which was increased from 3 to 12 for better model quality and to avoid steric clashes. The local quality of the model was assessed by predicted local distance difference test (pLDDT) scores as returned by AlphaFold. The confidence in the relative arrangement between NP and Z proteins was evaluated by predicted aligned errors (PAE), also as returned by AlphaFold. Other evaluations of the model quality and properties, including Predicted DockQ score (pDockQ) [72], protein-interface score (PI-score) [73], and biophysical properties of the interaction interface (using PI-score program), were calculated by AlphaPulldown.

### Mass photometry

Mass photometry was performed on a Refeyn OneMP mass photometer using the programs Acquire MP v2.5.0 and Discover MP v2.5.0 (Refeyn Ltd.) and PhotoMol. The workflow was published previously [74]. NP was measured in 50 mM Tris (HCl), 150 mM NaCl, 5 % Glycerol and a concentration of 25 nM. For the NP + RNA condition, NP at 500 nM was mixed with a single stranded 9mer (Table S3) at a 4 molar-excess to the NP monomer. NP + RNA was incubated at room temperature for approximately 5 min. The sample was diluted with 50 mM Tris (HCl), 150 mM NaCl, 5 % Glycerol to a concentration of 25 mM and subsequently measured.

### Experimental design and Statistical analysis

The determined masses, FWHM and AUC from nMS analysis were taken from at least 3 independent measurements and are shown together with the resulting standard deviation. Values for quantification are shown as mean from three independent measurements. The error bars represent the standard deviation. The shown standard deviation for K_d_ values are calculated according to the rules of Gaussian error propagation. HDX-MS experimental design and data analysis was evaluated according to HDX-MS community-recommendations [75]. The quench conditions were optimized for maximum sequence coverage. The back exchange was tested with a fully deuterated control. The labeling timepoints cover 3-4 orders of magnitude. The deuterium uptake differences between two states were statistically evaluated by using a two paired t-test with the p-value < 0.05. Peptides showing differences were only considered to be significant if passing the t-test and have additionally a difference higher than 0.561 deuteron, which is the variance across all replicates of all peptides.

## Acknowledgments

We thank Prof. H. Schlüter (UKE, University of Hamburg, Germany) for access to high-resolution mass spectrometer for the HDX experiments. We would like to thank Janine-Denise Kopicki for helpful advice of the K_D_ calculation. We thank Kay Grünewald for support and all current and past members of the Uetrecht and Rosenthal group for discussions and data interpretation. Part of this work was performed at the Multi-User CryoEM Facility at the Centre for Structural Systems Biology, Hamburg, supported by the Universität Hamburg and DFG grant numbers (INST 152/772-1|152/774-1|152/775-1|152/776-1|152/777-1 FUGG)

## Funding

Leibniz Center Infection Graduate School and Leibniz Interact (to MR, CU)

The Leibniz Institute of Virology was supported by the Free and Hanseatic City of Hamburg and the Federal Ministry of Health (to LS, CU)

German Federal Ministry of Education and Research 01KI2019 (to MR)

DFG RO 5954/2-1 (to MR)

DFG KO 5979/2-1 (to JK)

Leibniz Association’s Leibniz competition programme K72/2017 (to MR) and SAW-2014-HPI-4 grant (to CU)

Horizon 2020 ERC StG-2017 759661 (to CU)

BWFBG of the Free and Hanseatic City of Hamburg for an equipment grant (CU)

## Author contributions

Conceptualization: LS, MR, CU

Methodology: LS, HW, DY, DV, JK, MR, CU

Investigation: LS, HW, DY

Visualization: LS, DY

Supervision: JK, MR, CU

Funding acquisition: MR, CU

Writing—original draft: LS

Writing—review & editing: LS, DY, JK, MR, CU,

## Data and materials availability

The AlphaFold model of the N-Z dimer will be deposited in ModelArchive upon manuscript acceptance. (https://www.modelarchive.org/).

Mass spectrometry raw data and the analyzed HDX dataset will be deposited in the PRIDE database upon manuscript acceptance.

