## Supplementary information for "An RNA to rule them all: Critical steps in Lassa virus ribonucleoparticle assembly and recruitment"

Lennart Sänger *et al.*

**This PDF file includes:**

Supplementary Text
Figs. S1 to S10
Table S1 to S4
References (1 to 3)

**Fig. S1.**

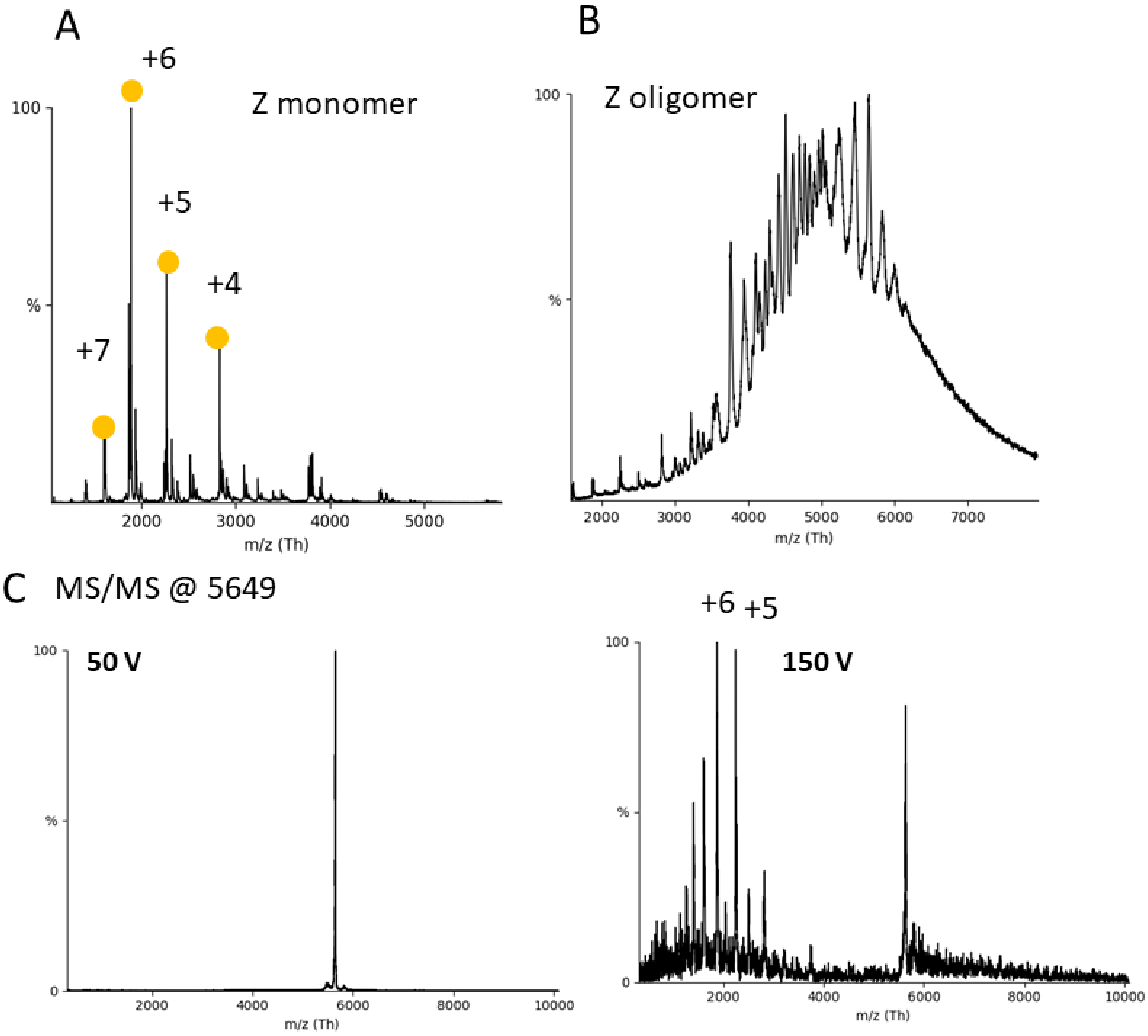


**nMS spectra of different oligomerization states of Z.** (A) The monomeric fraction of Z was measured with native MS in a 150 mM ammonium acetate buffer surrogate at pH 7.5. Capillary and cone voltage were constantly at 1450 V and 150 V, respectively. Acceleration voltage in the collision cell was at 50 V. Z appeared primarily in the monomeric state (11.30 +/-0.04 kDa) between 1000 and 4000 *m/z.* The charge states between +4 and +7 are depicted. Small fractions of dimers were traceable. (B) The oligomeric fraction of Z was measured with the same setting as described in A. Heterogenous peaks were measured between 4000 and 7000 *m/z*. (C) CID-MS/MS of one peak corresponding to a Z oligomer at 5649 *m/z* with low and high CV. Low *m/z* CID product at high CV corresponds to the mass of a Z monomer.

**Fig. S2.**

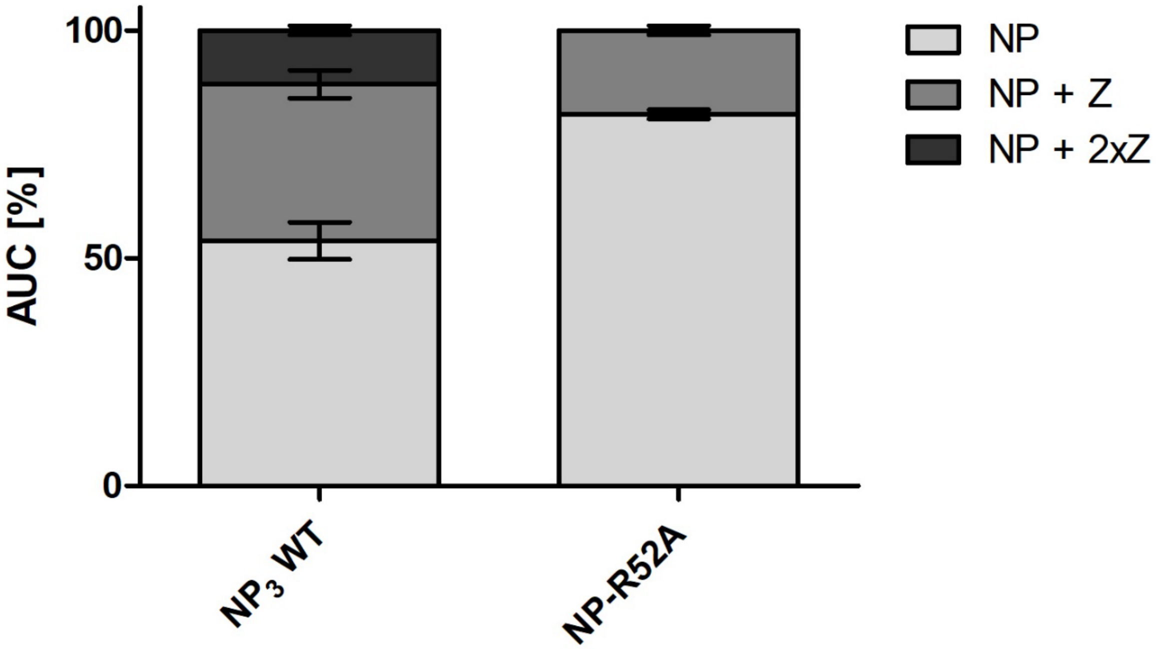


**Overall area under the curve (AUC) for the NP-Z mass species for NP WT and NP_R52A $K_D$ determination.** $NP_3$ WT and NP-R52A were incubated in a 1:3 (NP:Z) with a NP monomer concentration of 9 and 3 µM respectively. Samples were subjected to nMS. Capillary voltages were held at 1.2 kV, source temperature at 50°C and HCD voltage at 100 V. Resulting spectra from at least 3 independent measurements were deconvoluted to a zero-charge mass spectrum with Unidec and the AUC was determined for the respective mass species. The $K_D$ was determined after [1] taking into account that NP WT contains as a trimer 3 Z binding pockets and the mutant as a monomer only one Z binding pocket.

**Fig. S3.**

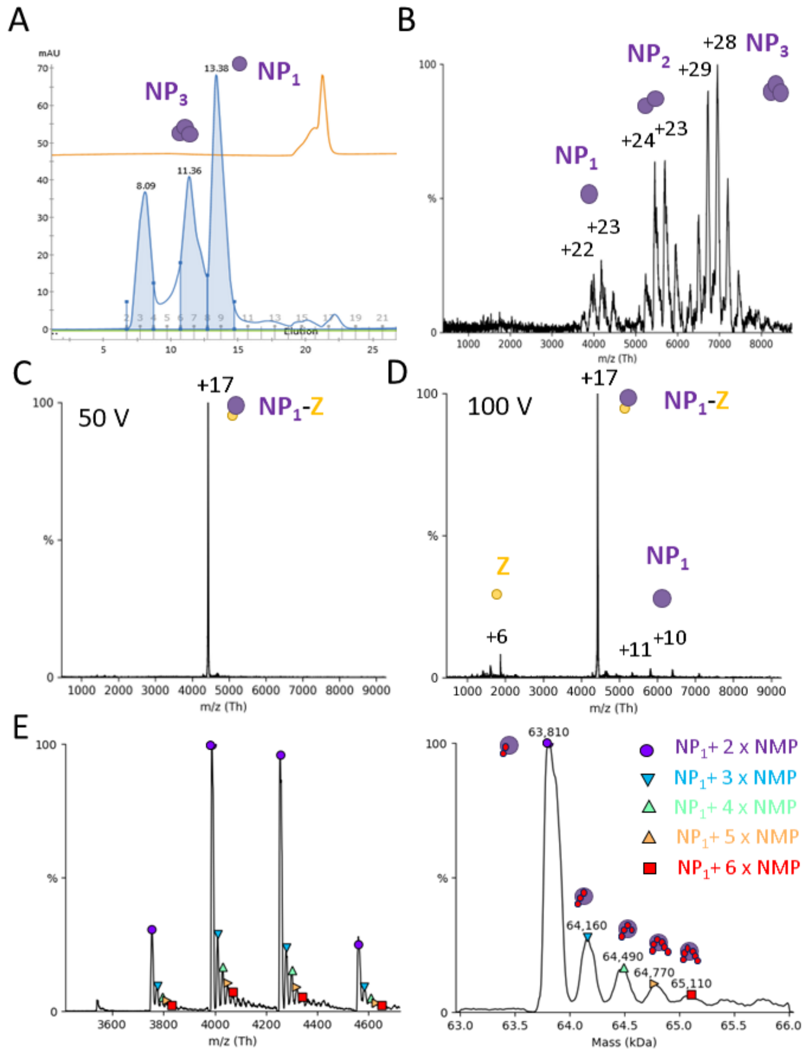


**Characterization of the NP trimerization mutant R52A.** (A) Size exclusion profile of R52A. The protein eluted in two fractions at 11.36 ml and 13.38 ml elution volume. nMS measurements of the peak at 13.38 revealed the monomeric character of the protein (Fig. 3A). nMS measurement of the peak at 11.36 ml elution volume is shown in (B). 150 mM ammonium acetate buffer surrogate at pH 7.5 was used for the measurement. Capillary and cone voltages were constantly at 1450 and 150 V, respectively. Acceleration voltage in the collision cell was at 50 V. The most abundant mass species was trimeric with the depicted +29 and +28 charge states. Between 4000 and 6000 *m/z* two peak series were visible with masses corresponding to NP dimers (+24 and +23 charge state) and NP monomers (+22 and +23 charge state). CID-MS/MS of one peak corresponding to a R52A-Z complex at 4431 *m/z* with low (C) and high (D) acceleration voltage. CID product at high acceleration voltage corresponds to a mass of a Z monomer and R52A. (E) Measurement of 4.1 µM NP R52A monomer fraction on a Orbitrap Q Exactive UHMR with high resolution, raw spectrum on the left revealing bound nucleotide monophosphates (NMPs) and deconvoluted spectrum on the right. 150 mM ammonium acetate solution at pH 7.5 was used for the measurement (see methods).

**Fig. S4.**

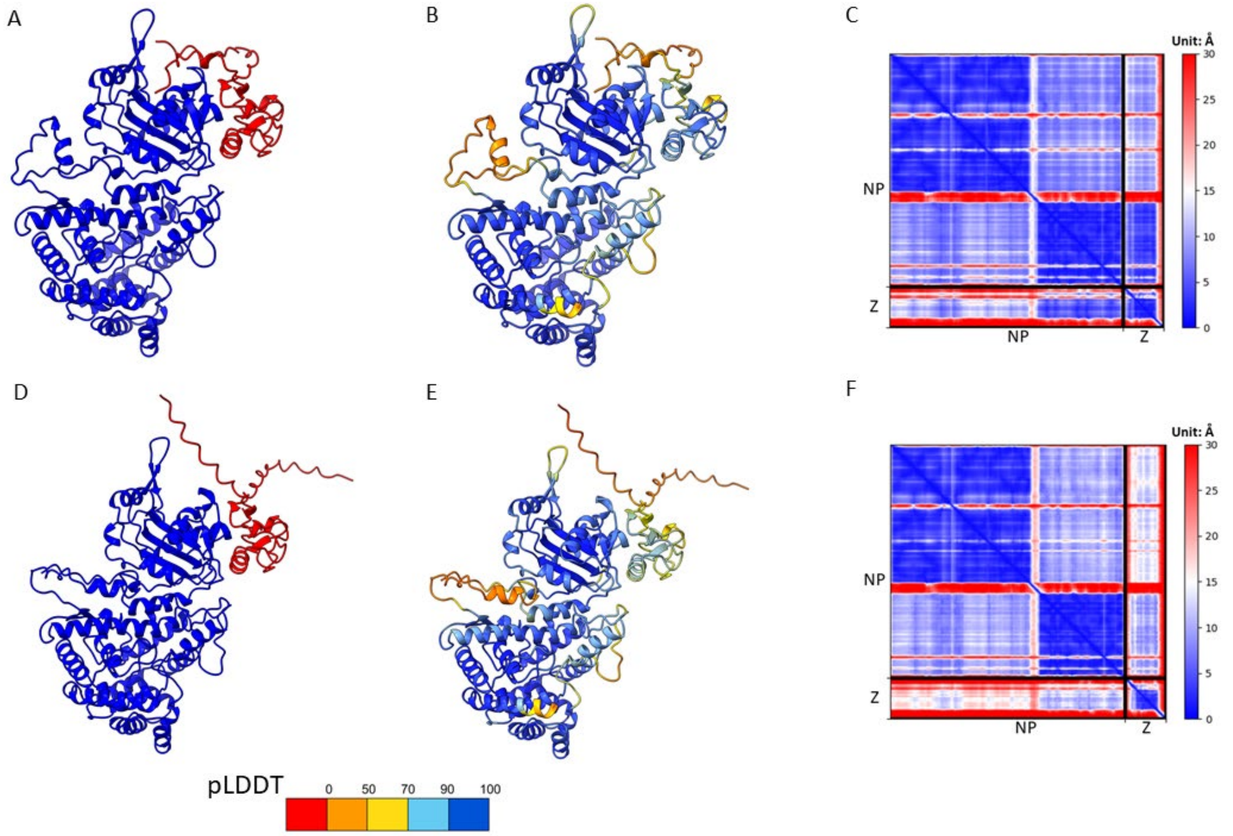


**Estimated quality of structural models for LASV NP and Z protein complex, built with the first AlphaFold Multimer version 2.1.0 and the latest AlphaFold Multimer version 2.3.0**. (A-C) NP-Z model predicted by AlphaFold Multimer version 2.1.0 colored by chains with NP in blue and Z protein in red; NP-Z model colored by each residue's predicted local distance difference test (pLDDT) score. A residue with a pLDDT score that is greater than 90 (dark blue) indicates high estimated accuracy of the position of its backbone and sidechain rotamers. A pLDDT score above 70 (light blue) suggests that the prediction of the backbone is confident; the predicted alignment error (PAE) plot of the NP-Z model shows the estimated expected distance error in Å. Each position (i,j) in the matrix is filled with the expected distance error in the residue i's position if the model and true (unknown) structure are aligned on residue j. All the PAE scores are reported by AlphaFold and blue means low errors in this plot. (D-F) NP-Z model predicted by AlphaFold Multimer version 2.3.0 colored by chains; NP-Z model colored by each residue's pLDDT score; the PAE plot of the NP-Z model. The two versions of AlphaFold Multimer use different sets of neural network weights and the consistency of the predictions additionally supports the models. Version 2.1.0 also exhibits higher pLDDT scores in the Z protein. These high scores appear reliable as the Z protein model agrees with the Z protein crystal structure.

**Fig. S5.**

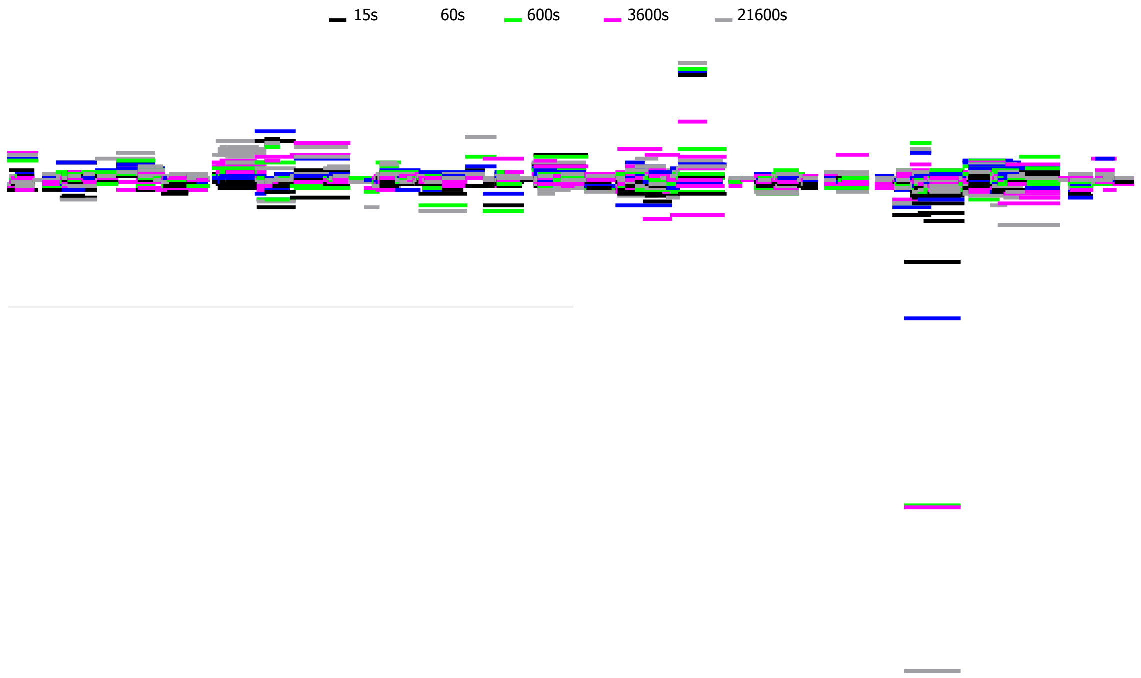


**Woodplot of HDX-MS data of NP together with Z.** HDX-MS dataset was analyzed with HDExaminer Version 3.3. Differences of deuterium uptake of NP peptides in presence of Z compared to an NP only control. Colors represent the different timepoints.

**Fig. S6.**

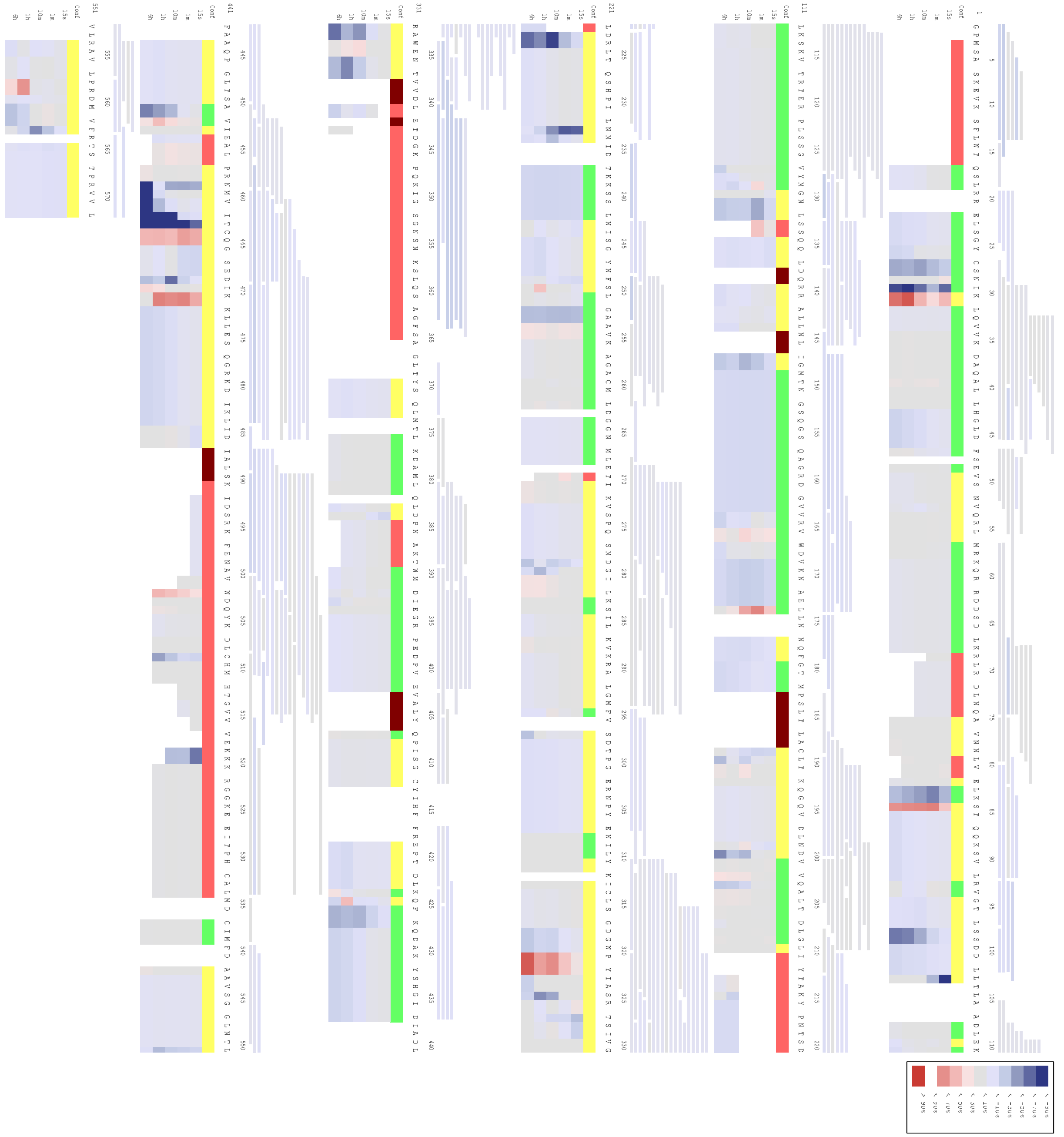

**HDX-MS analysis of NP and NP together with Z.** Heatmap of the HDX-MS analysis of NP and NP in presence of Z. HDX-MS dataset was analyzed with HDExaminer Version 3.3. The bars below the sequence represent heat bars of differences between the two states (NP and NP together with Z) based on the atomic range. The differences are colored from blue (protection) to red (exposure). Grey represents no differences between both states. The confidence of every residue is represented by the top bar where red means low confidence, yellow means medium confidence, and green means high confidence calculated by HDExaminer

**Fig. S7.**

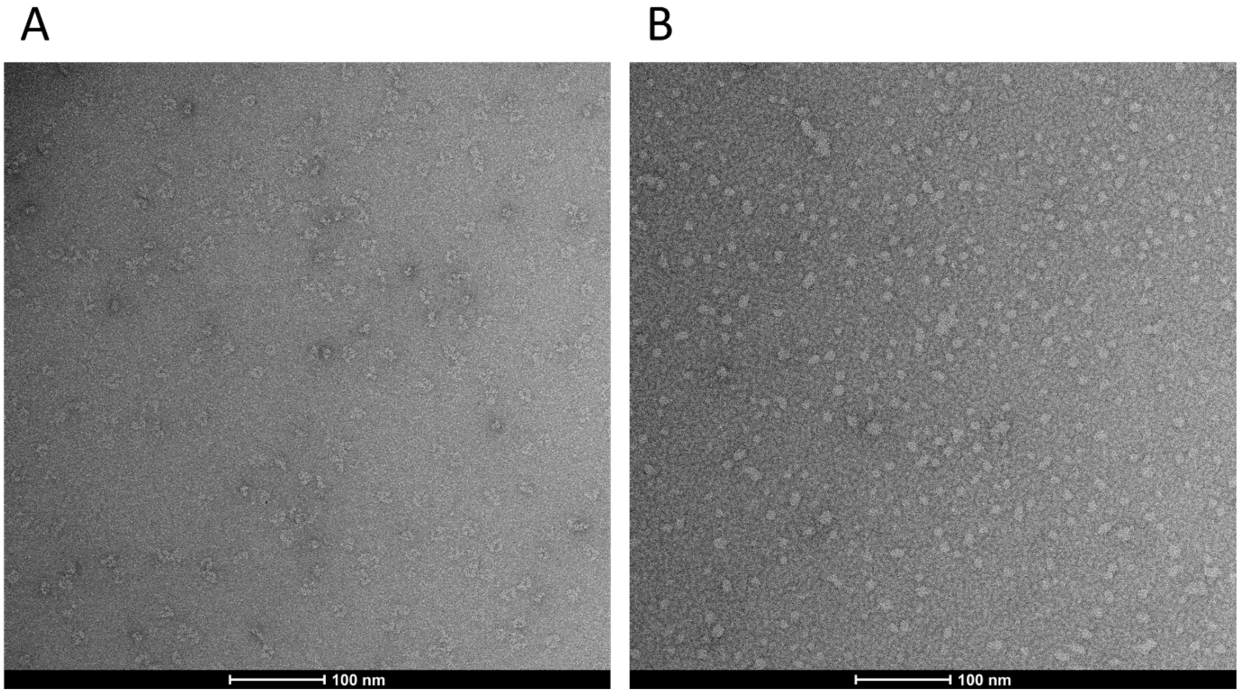


**Negative-stain electron microscopy of NP.** (A) NP at 0.01 mg/mL was applied to glow-discharged carbon-coated copper grids and stained with uranyl acetate immediately before imaging. Images were collected with a transmission electron microscopy. (B) For the NP + RNA condition, NP at 0.01 mg/mL was mixed with a single-stranded 25mer at a 1:2 molar ratio (NP:RNA). NP and RNA was incubated at room temperature for approximately 15 min and imaged as the no RNA condition.

**Fig. S8.**

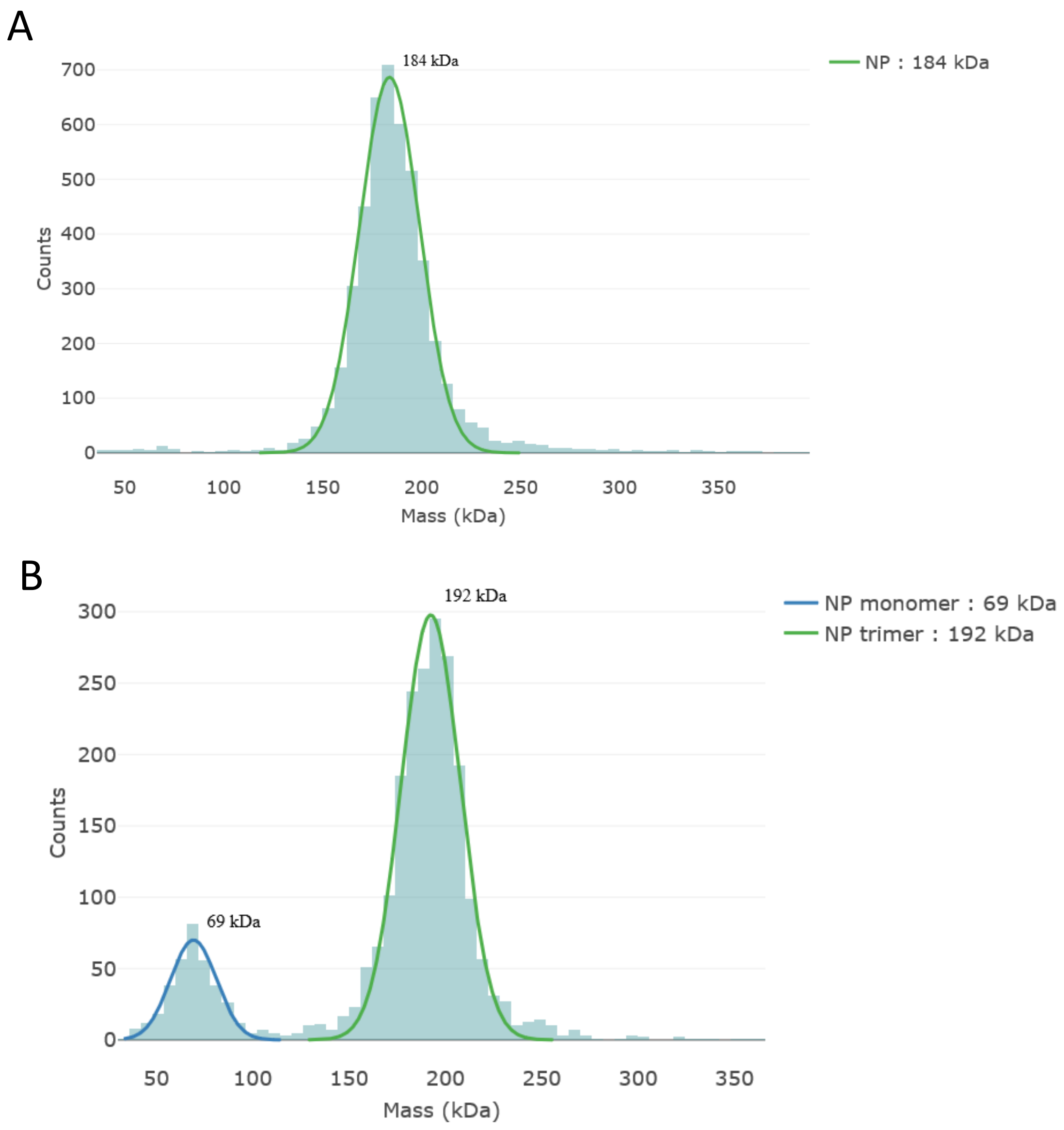


**Mass photometry of NP in presence or absence of RNA.** (A) Mass photometry was performed on a Refeyn OneMP mass photometer. NP was measured in a concentration of 25 nM in a physiological buffer condition. 4291 counts for NP were detected which represents 91 % of the total counts. The mass of NP was determined with 184 kDa and a sigma of 15 kDa. (B) For the NP + RNA condition, NP at 500 nM was mixed with a single stranded 9mer (Table S3) in a 4 molar-excess to the NP monomer. The NP + RNA sample was diluted to an NP concentration of 25 nM and subsequently measured. 1880 counts for a mass of 192 kDa (sigma 15 kDa) were detected, which represents 78 % of the total counts. The slightly higher mass of 192 kDa could indicate the $NP_3$-RNA intermediate state which we observed in our nMS experiments. Additionally, 344 counts for a mass of 69 kDa (sigma 12 kDa) were detected representing 14 % of the total counts.

**Fig. S9.**

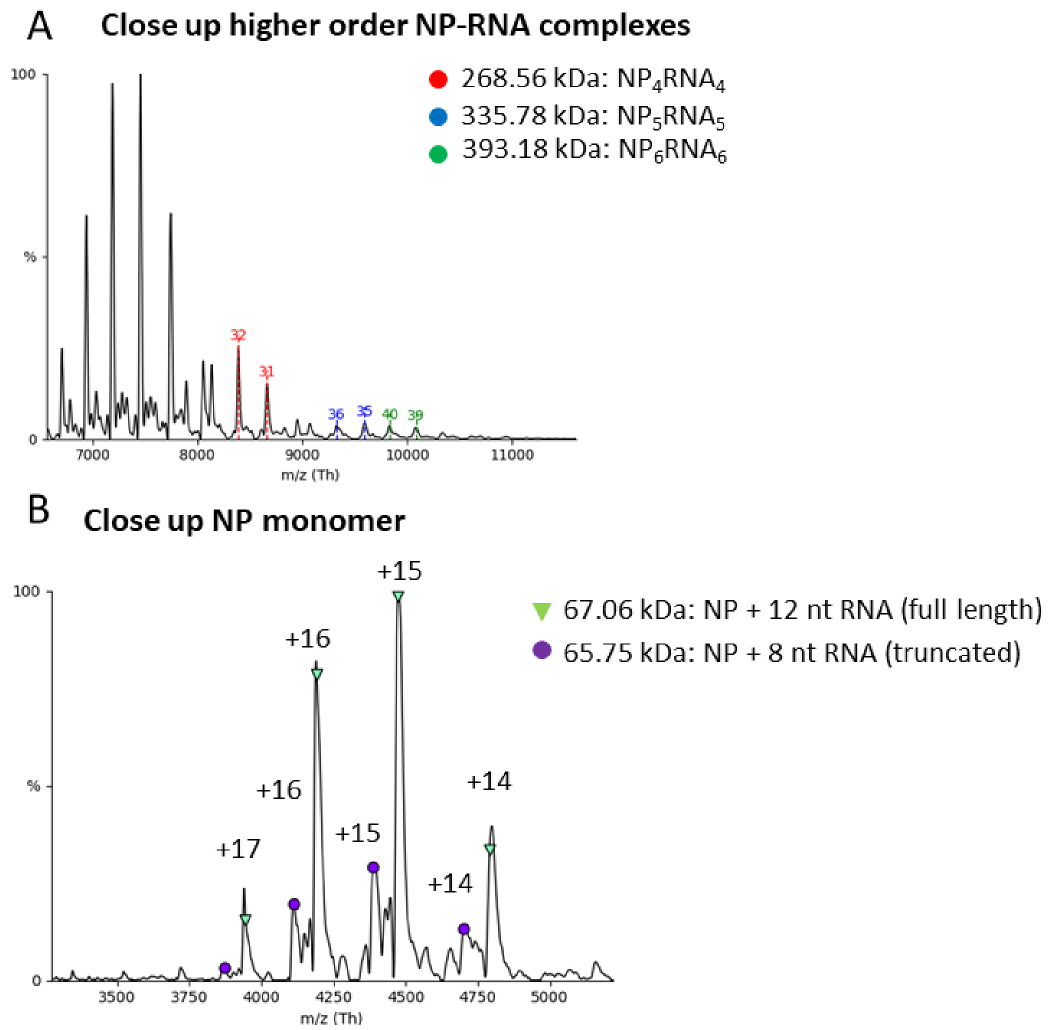


**nMS of NP-RNA species.** (A) NP was measured together with a 12 nt RNA in a 1:2 ratio (NP:RNA). Shown are spectra at a late timepoint of the reaction. Small fractions of higher order NP-RNA assemblies appear between 9000 and 11000 *m/z*. Masses corresponding to $NP_5RNA_5$ 335.78 kDa (FWHM: 0.90 kDa) and $NP_6RNA_6$ 393.18 kDa (FWHM: 1.12 kDa). Charge states of 36+, 35+ and 40+, 39+ are indicated. (B) Close up into the mass range of NP monomers. The main monomer-RNA species is indicated in green with a corresponding mass of 67.06 +/- 0.05 kDa. Smaller amounts of NP monomers with truncated bound RNAs are indicated in purple with a corresponding mass of 65.75 kDa (FWHM: 0.21 kDa). The mass difference is about 4 nucleotides.

**Fig. S10.**

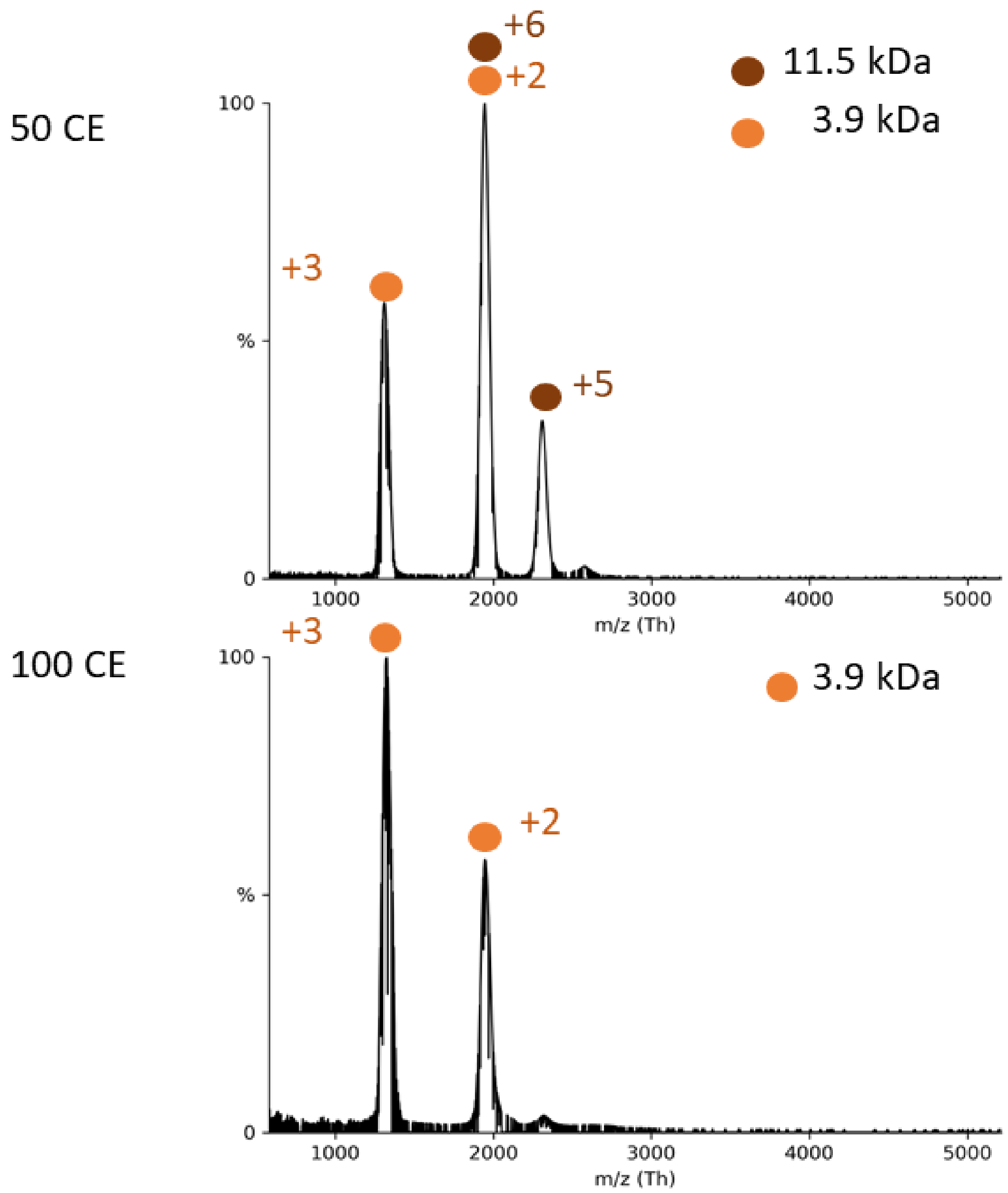


**nMS spectra of 9 nts RNA.** (A) 25 µM of the 12 nt RNA was measured at different collision energies. The theoretical mass of this RNA is 3.78 kDa. At 50 collisional energy two mass species were identified in the range between 1000 and 3000 *m/z*. Charge states of 3+, 2+ and 6+, 5+ are indicated. Masses correspond to 3.9 kDa and 11.5 kDa indicating single and a trimeric RNA structure. Only one peak series was observable at 100 CE corresponding to a mass of a single RNA (3.9 kDa).

**Table S1.**

NP-Z model quality assessments and interaction interface properties calculated by AlphaPulldown package. iPTM and iPTM+pTM scores are reported by AlphaFold. Both scores range between 0 and 1 with 1 being the best. pDockQ score is calculated using the formula given by [2]. pDockQ evaluates the quality of the interface and it ranges between 0.018 and 0.742 and a higher pDockQ score indicates better quality at the interface. The rest of the rows are evaluations reported by PI-score pipeline [3]. PI-score is a binary classifier that will assign a class label to a protein complex structure. A positive PI-score classifies the complex as a "native-like" fit whereas a negative PI-score indicates otherwise. Num_intf_residues: number of residues at the interface. Polar: number of polar residues (Ser, Thr, Asn, Gln, His and Tyr) at the interface. Hydrophobic: number of hydrophobic residues (Ala, Leu, Ile, Val, Phe, Trp, Cys, Met) at the interface. Charged: number of charged residues (Asp, Glu, Lys, Arg) at the interface. contact_pairs: number of atomic contacts between the interface residues. sc: geometric shape complementarity of protein-protein interfaces. sc ranges between 0 and 1 and sc=1 indicates two proteins mesh precisely. hb: number of hydrogen-bonds in the interface. sb: number of salt-bridges at the interface. int_solv_en: interface solvation energy. int_area: interface surface area that will be inaccessible to solvent upon the interface formation.

| | **AlphaFold_Multimer_V2.1.0** | **AlphaFold_Multimer_V2.3.0** |
|---|---|---|
| **Num_intf_residues** | 41 | 30 |
| **Polar** | 0.22 | 0.23 |
| **Hydrophobic** | 0.415 | 0.433 |
| **Charged** | 0.22 | 0.2 |
| **contact_pairs** | 41 | 33 |
| **sc** | 0.56 | 0.61 |
| **hb** | 7 | 6 |
| **sb** | 4 | 0 |
| **int_solv_en** | -11.25 | -8.1 |
| **int_area** | 1373.37 | 782.44 |
| **pi_score** | 0.77 | 1.27 |
| **iPTM+pTM** | 0.759 | 0.600 |
| **pDockQ** | 0.489 | 0.390 |
| **iPTM** | 0.748 | 0.559 |

**Table S2.**

| Data Set | unbound | 50 µM Z |
|---|---|---|
| **HDX reaction details** | 40 mM Tris, pH 7.5, 150 mM NaCl, 25°C | |
| **HDX time course (min)** | 0.25, 1, 10, 60, 360 | |
| **HDX control samples** | Fully deuterated control, labeled for 24 h, buffer 40 mM Tris, 6 M urea, pH 7.5 | |
| **Back-exchange (mean)** | 28.442 % +/- 8.537 % | |
| **# of Peptides** | 285 | 282 |
| **Sequence coverage** | 97.90% | 97.90% |
| **Average peptide length / Redundancy** | 13.80 / 6.89 | 13.71 / 6.77 |
| **Replicates** | 3 (technical) | 3 (technical) |
| **Repeatability (average standard deviation)** | 0.0716 | 0.0659 |
| **Significant differences in HDX (delta HDX > X D)** | t-test 95 % confidence interval and delta D < 0.5616 (variance across all peptides) | |

**Table S3.**

Sequences of RNA used in this study

| RNA | Sequence (5’-3’) |
|---|---|
| **9 nt RNA** | UAGGAAUCU |
| **12 nt RNA** | ACACAAAGACCC |
| **25 nt RNA** | GCCUAGGAUCCACUGUGCGUGUUGU |

**Table S4.**

**Experimental masses and FWHM for all proteins of this study obtained by nMS.**
Experimental masses ($M_{exp}$) of the different protein species in absence or presence of different ligands are determined from three independent measurements are listed together with standard deviation s along with the theoretically calculated molecular weight (Mass) based on the peptide chain. Additionally, the average full width of the peak at half maximum (FWHM) is listed.

| Mass species | Mass /Da | $M_{exp}$ /Da | s ($M_{exp}$) /Da | FWHM /Da |
|---|---|---|---|---|
| $NP_3$ | 189168 | 189557 | 25 | 784 |
| Z | 11187 | 11347 | 38 | 12 |
| R52A | 62971 | 63900 | 82 | 433 |
| R52A-Z | 74158 | 75473 | 232 | 519 |
| $NP_1$-$RNA12_1$ | 66840 | 67033 | 47 | 313 |
| $NP_2$-$RNA12_2$ | 133680 | 134033 | 47 | 509 |
| $NP_3$-$RNA12_3$ | 200520 | 201167 | 125 | 638 |
| $NP_3$-$RNA12_3$ | 192952 | 193633 | 94 | 583 |
| $NP_4$-$RNA12_4$ | 267360 | 268250 | 50 | 709 |
| $NP_5$-$RNA12_5$ | 334200 | 335780 | n/d | 897 |
| $NP_6$-$RNA12_6$ | 401040 | 393181 | n/d | 1116 |
| $NP_3$-$RNA9_1$ | 192007 | 193000 | 82 | 1077 |
| $NP_3$-$Z_1$ | 200355 | 201000 | 82 | 1258 |
| $NP_3$-$Z_2$ | 211542 | 212367 | 125 | 330 |
| $NP_3$-$Z_3$ | 222729 | 223867 | 170 | 319 |
| $NP_1$-$RNA9_1$ | 65895 | 66085 | 22 | 784 |
| $NP_1$-$Z_1$-$RNA9_1$ | 77082 | 77377 | 56 | 12 |

1. Kopicki, J.-D., et al., *Opening opportunities for Kd determination and screening of MHC peptide complexes.* Communications Biology, 2022. **5**(1).
2. Bryant, P., G. Pozzati, and A. Elofsson, *Improved prediction of protein-protein interactions using AlphaFold2.* Nature communications, 2022. **13**(1): p. 1-11.
3. Malhotra, S., et al., *Assessment of protein–protein interfaces in cryo-EM derived assemblies.* Nature communications, 2021. **12**(1): p. 1-12.
